# How the forest interacts with the trees: Multiscale shape integration explains global and local processing

**DOI:** 10.1101/777110

**Authors:** Georgin Jacob, S. P. Arun

## Abstract

Hierarchical stimuli (such as a circle made of diamonds) have been widely used to study global and local processing. Two classic phenomena have been observed using these stimuli: the global advantage effect (that we identify the circle faster than the diamonds) and the incongruence effect (that we identify the circle faster when both global and local shapes are circles). Understanding them has been difficult because they occur during shape detection, where an unknown categorical judgement is made on an unknown feature representation.

Here we report two essential findings. First, these phenomena are present both in a general same-different task and a visual search task, suggesting that they may be intrinsic properties of the underlying representation. Second, in both tasks, responses were explained using linear models that combined multiscale shape differences and shape distinctiveness. Thus, global and local processing can be understood as properties of a systematic underlying feature representation.

## INTRODUCTION

Visual objects contain features at multiple spatial scales (Oliva and Schyns, 1997; Morrison and Schyns, 2001; Ullman et al., 2002). Our perception of global and local shape have been extensively investigated using hierarchical stimuli, which contain local elements arranged to form a global shape (Figure 1). Two classic phenomena have been observed using these stimuli (Navon, 1977; Kimchi, 1992). First, the global shape can be detected faster than the local shape; this is known as the global advantage effect. Second, the global shape can be detected faster in a congruent shape (e.g. circle made of circles) than in an incongruent shape (e.g. circle made of diamonds); this is known as the global-local incongruence effect. Subsequent studies have shown that these effects depend on the size, position, spacing and arrangement of the local shapes (Lamb and Robertson, 1990; Kimchi, 1992; Malinowski et al., 2002; Miller and Navon, 2002).

**Figure 1.**
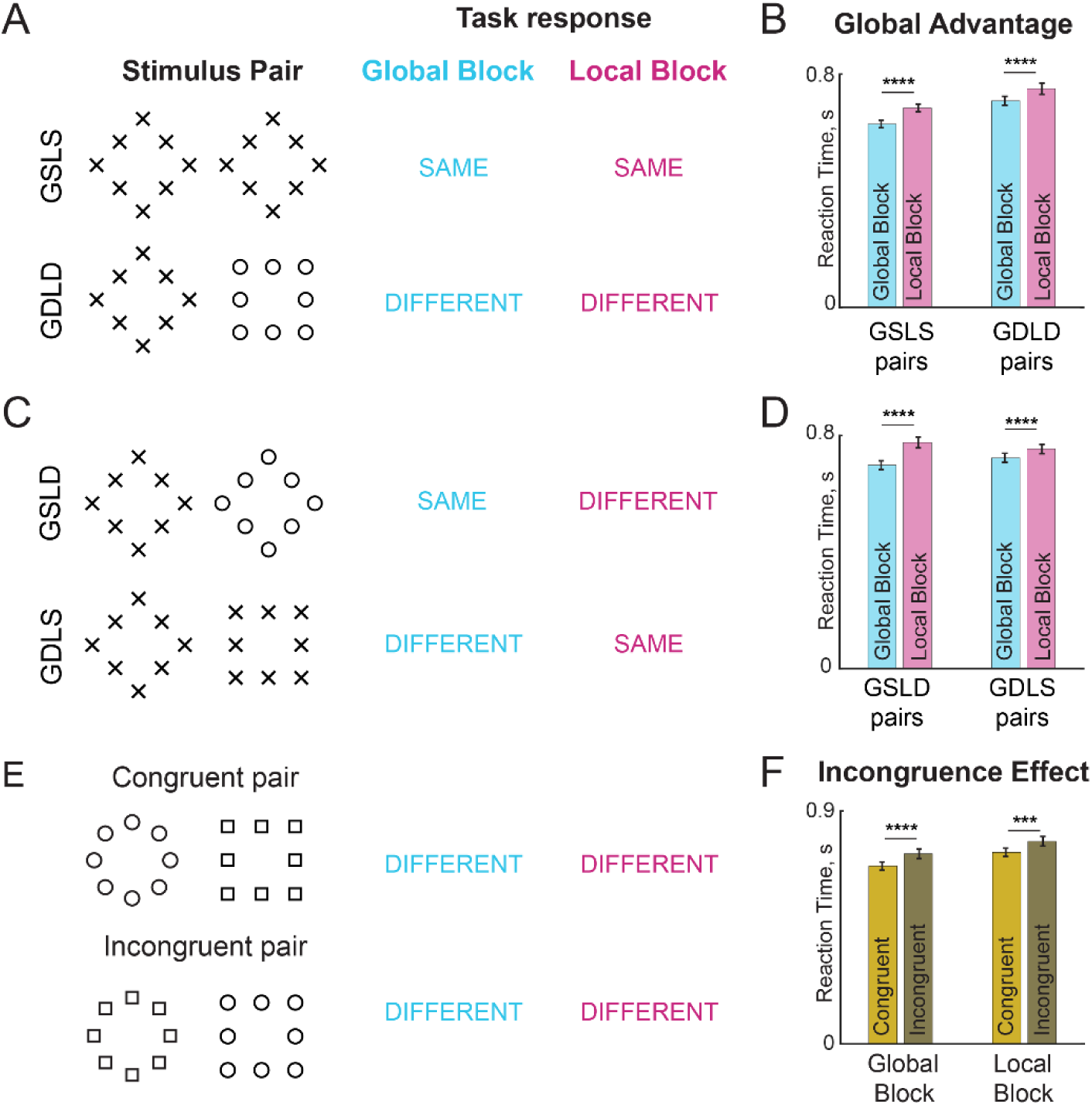
Same-different task for global-local processing. In the global block, subjects have to indicate whether a pair of images presented contain the same shape at the global level. Likewise in the local block, they have to make same-different judgments about the shape at the local level. Block order was counterbalanced across subjects. (A) Example image-pairs with identical correct responses in the global and local blocks. In the GSLS pairs, both images are identical i.e. have the same global shape and same local shape. In the GDLD pairs, the two images differ in both global shape and local shape. (B) Bar plot comparing average response times for GSLS and GDLD pairs. Error bars indicate s.e.m. across subjects. Asterisks indicate statistical significance assessed using an ANOVA on response times (**** is p < 0.00005). (C) Example image pairs that elicited opposite responses in the global and local blocks. In the GSLD pairs, the two images contain the same global shape but differ in local shape – thus the correct response is “SAME” in the global block but “DIFFERENT” in the local block. In the GDLS pairs, the two images contain the same local shape but differ in global shape, resulting again in opposite responses in the two blocks. (D) Same as B but for GSLD and GDLS pairs. (E) Example congruent and incongruent image pairs. Congruent image pairs comprised stimuli with the same shape at the global and local levels. In the incongruent image pairs, the global shape of one image matched the local shape of the other, and vice-versa. Thus each congruent image pair was exactly matched to an incongruent image pair. (F) Bar plot of average response times for congruent and incongruent image pairs. Asterisks indicate statistical significance using an ANOVA on response times (**** is p < 0.00005).

These global/local processing phenomena have since been extensively investigated for their neural basis as well as their application to a variety of disorders. Global and local processing are thought to be localized to the right and left hemispheres respectively (Fink et al., 1996; Han et al., 2002, 2004), and are mediated by brain oscillations at different frequencies (Romei et al., 2011; Liu and Luo, 2019). These phenomena have now been observed in a variety of other animals, especially during tasks that require speeded responses (Tanaka and Fujita, 2000; Cavoto and Cook, 2001; Pitteri et al., 2014; Avarguès-Weber et al., 2015). Global/local processing is impaired in a variety of clinical disorders (Bihrle et al., 1989; Robertson and Lamb, 1991; Slavin et al., 2002; Behrmann et al., 2006; Song and Hakoda, 2015), including those related to reading (Lachmann and Van Leeuwen, 2008; Franceschini et al., 2017). Finally, individual differences in global/local processing predict other aspects of object perception (Gerlach and Poirel, 2018; Gerlach and Starrfelt, 2018).

Despite these insights, we lack a deeper understanding of these phenomena for several reasons. First, they have only been observed during shape detection tasks, which involve two complex steps: a categorical response made over a complex underlying representation (Freedman and Miller, 2008; Mohan and Arun, 2012). It is therefore possible that these phenomena reflect the priorities of the categorical decision. Alternatively, they may reflect some intrinsic property of the underlying shape representation.

Second, these shape detection tasks, by their design, set up a response conflict for incongruent but not congruent stimuli. This is because the incongruent stimulus contains two different shapes at the global and local levels, each associated with a different response during the global and local blocks. By contrast there is no such conflict for congruent stimuli where the global and local shapes are identical. Thus, the incongruence effect might reflect the response conflicts associated with making opposite responses in the global and local blocks (Miller and Navon, 2002). Alternatively, again, it might reflect some intrinsic property of the underlying shape representation.

Third, it has long been appreciated that these phenomena depend on stimulus properties such as the size, position, spacing and arrangement of the local elements (Lamb and Robertson, 1990; Kimchi, 1992; Malinowski et al., 2002; Miller and Navon, 2002). Surprisingly, hierarchical stimuli themselves have never been studied from the perspective of feature integration i.e. how the global and local shapes combine. A deeper understanding of how hierarchical stimuli are organized in perception can elucidate how these stimulus properties affect global/local processing.

Thus, understanding the global advantage and incongruence effects will require reproducing them in simpler tasks, as well as understanding how global and local shape combine in the perception of hierarchical stimuli. This is not only a fundamental question but has clinical significance since deficits in global/local processing have been reported in a variety of disorders.

### Overview of this study

Here we addressed the above limitations as follows. First, we devised a simpler shape task which involves subjects indicating whether two shapes are the same or different at either the global or local level. This avoids any effects due to specific shapes but still involves categorization, albeit a more general one. Second, we devised a visual search task in which subjects had to report the location of an oddball target. This task avoids any categorical judgement and the accompanying response conflicts. It also does not involve any explicit manipulation of global vs local attention unlike the global/local processing tasks. If these phenomena are present in visual search, it would imply that they reflect properties of the underlying shape representation of hierarchical stimuli. If not, they must arise from the categorization process.

To understand how global and local shape combine in visual search, we asked how search difficulty for a target differing in both global and local shape from the distractors can be understood in terms of global and local shape differences. While search reaction time (RT) is the natural observation made during any search task, we have shown recently that its reciprocal (1/RT) is the more useful measure for understanding visual search (Arun, 2012; Pramod and Arun, 2014). The reciprocal of search time can be thought of as the dissimilarity between the target and distractors in visual search, and has the intuitive interpretation as the underlying salience signal that accumulates to threshold (Arun, 2012). Models based on 1/RT consistently outperform models based directly on search time (Vighneshvel and Arun, 2013; Pramod and Arun, 2014, 2016; Sunder and Arun, 2016). Further, using this measure, a variety of object attributes as well as top-down factors such as target preview have been found to combine linearly.

We performed two experiments. In Experiment 1, we replicated the global advantage and incongruence effects in a generic same-different task. We then show that image-by-image variations in response times can be explained by two factors: dissimilarity and distinctiveness. In Experiment 2, we show that these effects can be observed even when subjects perform visual search on the same stimuli. We also show that visual search for hierarchical stimuli can be accurately explained as a linear sum of global and local feature relations. Finally we show that the factors driving the same-different task responses are closely related to the visual search model.

## EXPERIMENT 1: SAME-DIFFERENT TASK

In most studies of global and local processing, subjects are required to indicate which of two target shapes they saw at the global or local levels (Navon, 1977; Kimchi, 1994). This approach severely limits the number of shapes that can be tested because of the combinatorial increase in the number of possible shape pairs. To overcome this limitation, we devised a same-different task in which subjects have to indicate whether two simultaneously presented shapes contain the same or different shape at the global or local level. Of particular interest to us were two questions: (1) Are the global advantage and incongruence effects observable in this more general shape detection task? (2) Do response times in this task systematically vary across stimuli and across the global and local blocks?

### METHODS

Here and in all experiments, subjects had normal or corrected-to-normal vision and gave written informed consent to an experimental protocol approved by the Institutional Human Ethics Committee of the Indian Institute of Science, Bangalore. Subjects were naive to the purpose of the experiment and received monetary compensation for their participation.

#### Subjects

Sixteen human subjects (11 male, aged 20-30 years) participated in this experiment. We chose this number of subjects based on previous studies of object categorization from our lab in which this sample size yielded consistent responses (Mohan and Arun, 2012).

#### Stimuli

We created hierarchical stimuli by placing eight local shapes uniformly along the perimeter of a global shape. All local shapes had the same area (0.77 squared degrees of visual angle), and all global shapes occupied an area that was 25 times larger. We used seven distinct shapes at the global and local levels to create 49 hierarchical stimuli (all stimuli can be seen in Figure 8). Stimuli were shown as white against a black background.

#### Procedure

Subjects were seated ∼60 cm from a computer monitor under the control of custom programs written in MATLAB with routines from PsychToolbox (Brainard, 1997). Subjects performed two blocks of the same-different task, corresponding to global or local shape matching. In both blocks, a pair of hierarchical shapes were shown to the subject and the subject had to respond if the shapes contained the same or different shape at a particular global/local level (key “Z” for same, “M” for different). Each block started with a practice block with eight trials involving hierarchical stimuli made of shapes that were not used in the main experiment. Subjects were given feedback after each trial during the practice block.

In all blocks, each trial started with a red fixation cross (measuring 0.6° by 0.6°) presented at the centre of the screen for 750 ms. This was followed by two hierarchical stimuli (with local elements measuring 0.6° along the longer dimension and longest dimension of global shapes are 3.8°) presented on either side of the fixation cross, separated by 8° from center to center. The position of each stimulus was jittered by ± 0.8° uniformly at random along the horizontal and vertical. These two stimuli were shown for 200 ms followed by a blank screen until the subject made a response, or until 5 seconds, whichever was sooner.

#### Stimulus pairs

To avoid any response bias, we selected stimulus pairs in each block such that the proportion of same- and different-responses were equal. Each block consisted of 588 stimulus pairs. These pairs were divided equally into four groups of 147 pairs (Figure 1A): (1) Pairs with global shape same, local shape same (GSLS, i.e. identical shapes); (2) Pairs with global shape same but local different (GSLD); (3) Pairs with global different but local same (GDLS) and (4) Pairs with both global and local shape different (GDLD). Since there were different number of total possible pairs in each category we selected pairs as follows: for GSLS pairs, there are 49 unique stimuli and therefore 49 pairs, so we repeated each pair three times to obtain 147 pairs. For GSLD and GDLS pairs, there are 147 unique pairs, so each pair was used exactly once. For GDLD pairs, there are 441 possible pairs, so we selected 147 pairs which consisted of 21 congruent pairs (i.e. each stimulus containing identical global and local shapes), 21 incongruent pairs (in which global shape of one stimulus was the local shape of the other, and vice-versa), and 105 randomly chosen other pairs. The full set of 588 stimulus pairs were fixed across all subjects. Each stimulus pair was shown twice. Thus each block consisted of 588 x 2 = 1176 trials. Error trials were repeated after a random number of other trials.

We removed inordinately long or short response times for each image pair using an automatic outlier detection procedure (*isoutlier* function, MATLAB 2018). We pooled the reaction times across subjects for each image pair, and all response times greater than three scaled median absolute deviations away from the median were removed. In practice this procedure removed ∼8% of the total responses.

#### Estimating data reliability

To estimate an upper limit on the performance of any model, we reasoned that the performance of any model cannot exceed the reliability of the data itself. To estimate the reliability of the data, we first calculated the average correlation between two halves of the data. However, doing so underestimates the true reliability since the correlation is based on two halves of the data rather than the entire dataset. To estimate this true reliability we applied a Spearman-Brown correction on the split-half correlation. This Spearman-Brown corrected correlation (*rc*) is given by *rc* = 2*r*/(1+*r*) where *r* is the correlation between the two halves. This data reliability is denoted as *rc* throughout the text to distinguish it from the standard Pearson’s correlation coefficient (denoted as *r*).

## RESULTS

Here, subject performed a same-different task in which they reported whether a pair of hierarchical stimuli contained the same/different shape at the global level or at the local level in separate blocks. We grouped the image pairs into four distinct types based on whether the shapes were same/different at the global/local levels. The first type comprised pairs with no difference at the global or local levels, i.e. identical images, denoted by GSLS (Figure 1A, top row). The second type comprised pairs in which both global and local shape were different, denoted by GDLD (Figure 1A, bottom row). These two were pairs elicited identical responses in the global and local blocks. The third type comprised pairs with the same global shape but different local shapes, denoted by GSLD (Figure 1C, top row). The fourth type comprised pairs differing in global shape but with identical local shapes, denoted by GDLS (Figure 1C, bottom row). These two were pairs that elicited opposite responses in the global and local blocks. Since both blocks consisted of identical image pairs, the responses in the two blocks are directly comparable and matched for image content.

### Is there a global advantage in the same-different task?

Subjects were highly accurate in the task overall, but were more accurate in the global block (mean & std of accuracy across subjects: 91% ± 4% in the global block; 88%±7% in the local block, p < 0.05, sign-rank test on subject-wise accuracy in the two blocks). They were also significantly faster in the global block (mean & std of response times across subjects: 702 ± 55.7 ms in the global block; 752 ± 66.7 ms in the local block; p < 0.005, sign-rank test on subject-wise average RTs in the two blocks). This pattern was true both for image pairs that elicited identical responses in the two blocks (GSLS & GDLD pairs; Figure 1B) as well as for those that elicited opposite responses (GDLS & GSLD pairs; Figure 1C). Thus, subjects were faster and more accurate in the global block across all image pairs, reflecting a robust global advantage.

### Is there an incongruence effect in the same-different task?

Next we asked whether the incongruence effect can be observed in the same-different task. To this end we compared the average RT for GDLD image pairs in which the two images were either both congruent or both incongruent (Figure 1E). Subjects responded significantly faster to congruent compared to incongruent pairs (Figure 1F). To assess the statistical significance of these effects, we performed an ANOVA on the response times with subject (16 levels), block (2 levels), congruence (2 levels) and image pair (21 levels) as factors. This revealed a significant main effect of congruence (p < 0.00005), but also main effects of subject and block (p < 0.00005 in all cases), as well as significant interaction effects (p < 0.00005, between subjects and blocks; all other effects were p > 0.05). We conclude that there is a robust incongruence effect in both the global and local blocks.

### Do responses in the same-different task vary systematically across image pairs?

Having established that subjects show a global advantage and incongruence effects in the same-different task, we wondered whether there were any other systematic variations in response times across image pairs. Specifically, we asked whether image pairs that evoked fast responses in one group of subjects would also elicit a fast response in another group of subjects. This was indeed the case: we found a significant correlation between the average response times of the first and second half of all subjects in both the global block (r = 0.74, p < 0.00005; Figure 2A) and the local block (r = 0.75, p < 0.000005; Figure 2B). This correlation was present in all four image types as well in both blocks (Figure 2).

**Figure 2.**
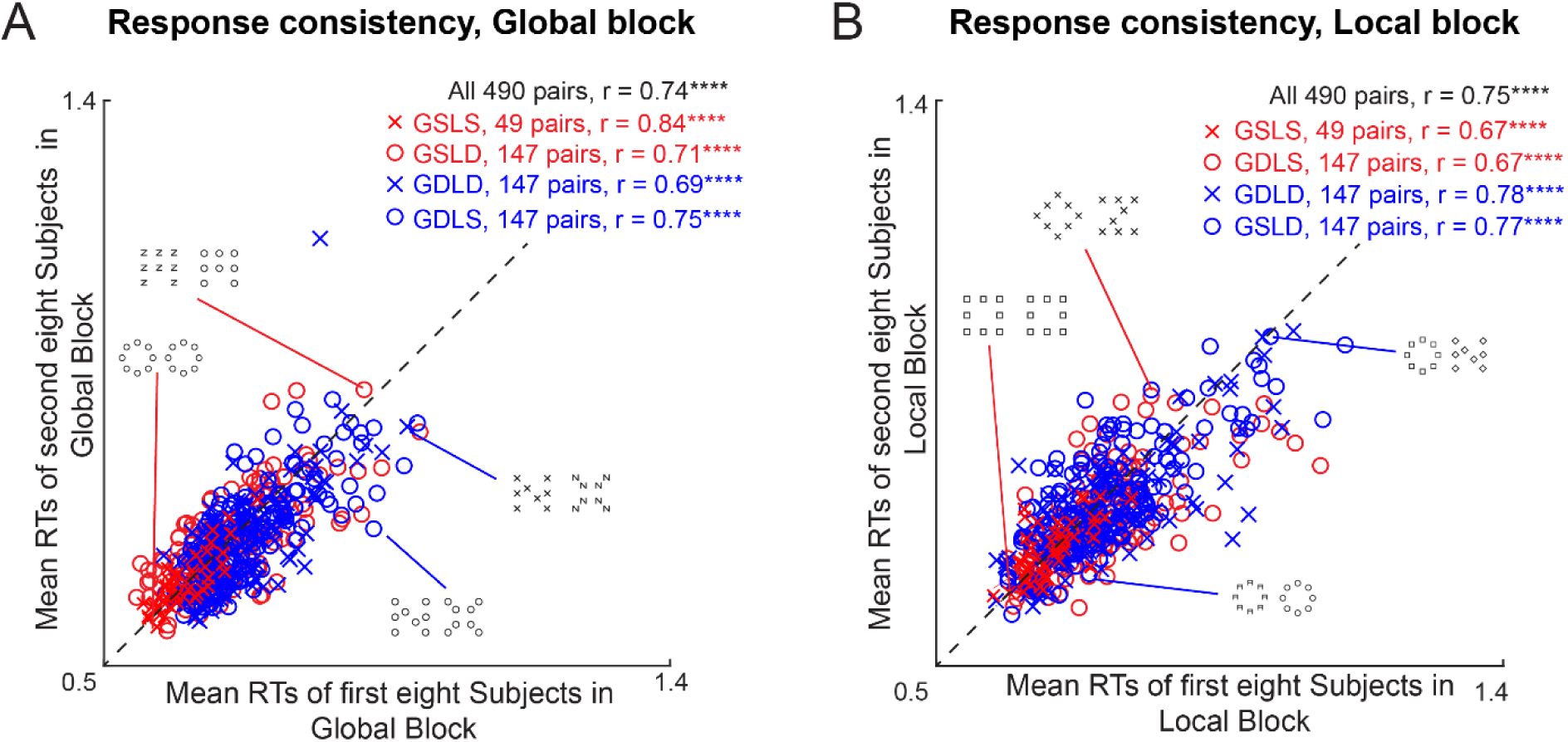
Consistency of response times in the same-different task. (A) Average response times for one half of the subjects in the global block of the same-different task plotted against those of the other half. Asterisks indicate statistical significance (* is p < 0.05, ** is p < 0.005 etc). (B) Same as (A) but for the local block.

### Are responses in the global and local block related?

Having established that response times are systematic within each block, we next investigated how responses in the global and local block are related for the same image pairs presented in both blocks. First, we compared responses to image pairs that elicit identical responses in both blocks. These are the GSLS pairs (which elicit a SAME response in both blocks) and GDLD pairs (that elicit a DIFFERENT response in both blocks). This revealed a positive but not significant correlation between the responses to the GSLS pairs in both blocks (r = 0.15, p = 0.32 across 49 image pairs; Figure 3A). By contrast the responses to the GDLD pairs, which were many more in number (n = 147), showed a significant positive correlation between the global and local blocks (r = 0.24, p < 0.005; Figure 3A). Second, we compared image pairs that elicited opposite responses in the global and local blocks, namely the GSLD and GDLS pairs. This revealed a significant negative correlation in both cases (r = −0.20, p < 0.05 for GSLD pairs, r = −0.23, p < 0.0005 for GDLS pairs; Figure 3B). Thus, image pairs that are hard to categorize as SAME are easier to categorize as DIFFERENT.

**Figure 3.**
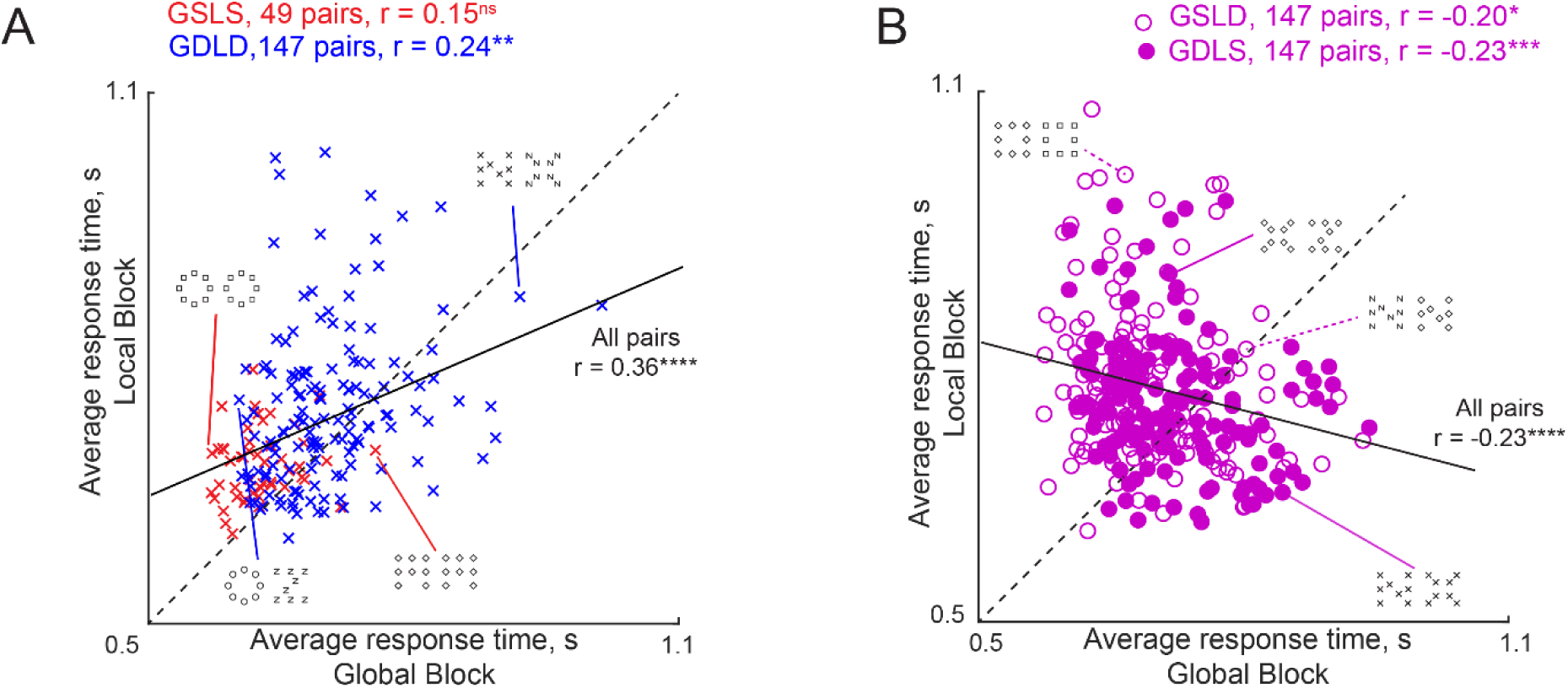
Responses to hierarchical stimuli in global and local blocks. (A) Average response times in the local block plotted against the global block, for image pairs with identical responses in the global and local blocks. These are the GSLS pairs (red crosses, n = 49) which elicited the “SAME” response in both blocks, and the GDLD pairs (blue crosses, n = 147) which elicited the “DIFFERENT” responses in both blocks. (B) Average response times in the local block plotted against the global block, for image pairs with opposite responses in the global and local blocks. These are the GSLD pairs (open circles, n = 147) which elicit the “SAME” response in the global block but the “DIFFERENT” response in the local block, and the GDLS pairs (filled circles, n = 147) which likewise elicit opposite responses in the two blocks.

Note that in all cases, the correlation between responses in the global and local blocks were relatively small (only r = ∼0.2; Figure 3) compared to the consistency of the responses within each block (split-half correlation = 0.75 in the global block; 0.74 in the local block; p < 0.00005 for both the conditions; Figure 2). These low correlations suggest that responses in the global and local blocks are qualitatively different.

### What factors influence response times in the same-different task?

So far we have shown that the global advantage and incongruence effects are present in a same-different task, and that response times vary systematically in each block across image pairs. However these findings do not explain why some image pairs elicit slower responses than others (Figure 2).

Consider a schematic of perceptual space depicted in Figure 4A. We hypothesized that the response time for an image pair in the global block could depend on two factors. The first factor is the dissimilarity between the two images. If the two images have the same global shape (thus requiring a “SAME” response), then the response time would be proportionally longer as the local shapes become more dissimilar. By contrast, if two images differ in global shape (thus requiring a “DIFFERENT” response), then the response time would be shorter if the two images are more dissimilar (Figure 4B). Thus, shape dissimilarity between the two images can have opposite effects on response time depending on whether the response is same or different. The second factor is the distinctiveness of the images relative to all other images. We reasoned that a shape that is distinct from all other shapes should evoke a faster response since there are fewer competing stimuli in its vicinity. This factor is required to explain systematic variation in response times for identical images (e.g. GSLS pairs) where the first factor (dissimilarity) plays no role. But more generally, distinctiveness could play a role even when both images are different. Below we describe how distinctiveness and dissimilarity can be used to predict response time variations in the same-different task.

**Figure 4.**
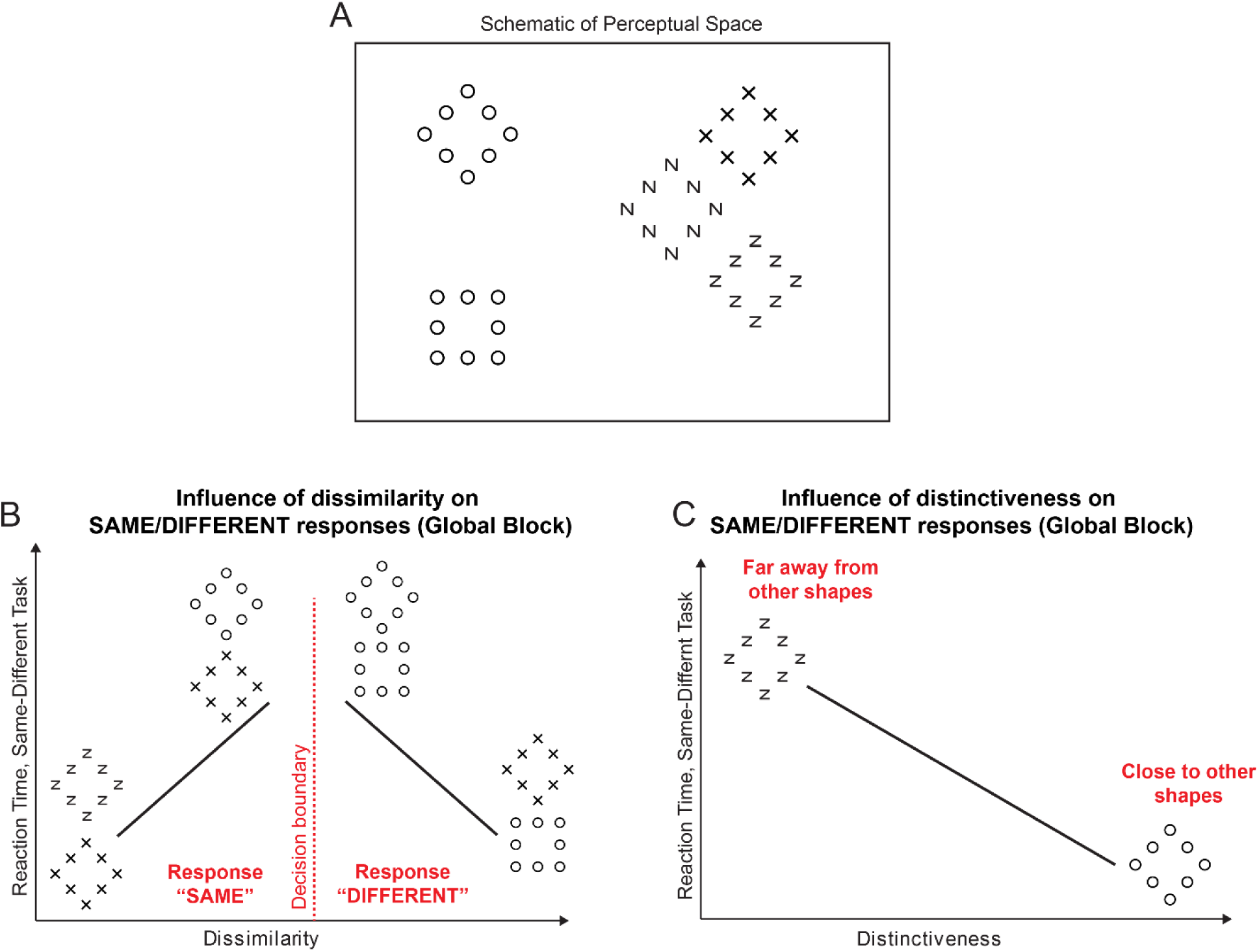
Understanding same-different responses. (A) To elucidate how same-different responses are related to the underlying perceptual space, consider a perceptual space consisting of many hierarchical stimuli. In this space, nearby stimuli are perceptually similar. (B) We hypothesized that subjects make “SAME” or “DIFFERENT” responses to an image pair driven by the dissimilarity between the two images. In the global block, when two images have the same global shape, we predict that response times are longer when the two images are more dissimilar. Thus, two diamonds made using Xs and Zs evoke a faster response than two diamonds made of circles or Xs, because the latter pair is more dissimilar than the former. By contrast, when two images differ in global shape, responses are faster when they are more dissimilar. (C) We also hypothesized that shapes that are more distinct i.e. far away from other shapes will elicit faster responses because there are no surrounding distractors. Thus, the diamond made of circles, which is far away from all other stimuli in panel A, will elicit a faster response than a diamond made of Zs.

### Effect of distinctiveness on same-different responses in the global block

How do we estimate distinctiveness? We reasoned that if distinctiveness was the only influence on response time to identical images, then images that elicited fast responses must be more distinctive than those that elicit slow responses. We therefore took the reciprocal of the average response time for each GSLS pair (across trials and subjects) as a measure of distinctiveness for that image. The estimated distinctiveness for the hierarchical stimuli in the global block is depicted in Figure 5A. It can be seen that shapes with a global circle (“O”) are more distinctive than shapes containing the global shape “A”. In other words, subjects responded faster when they saw these shapes.

**Figure 5.**
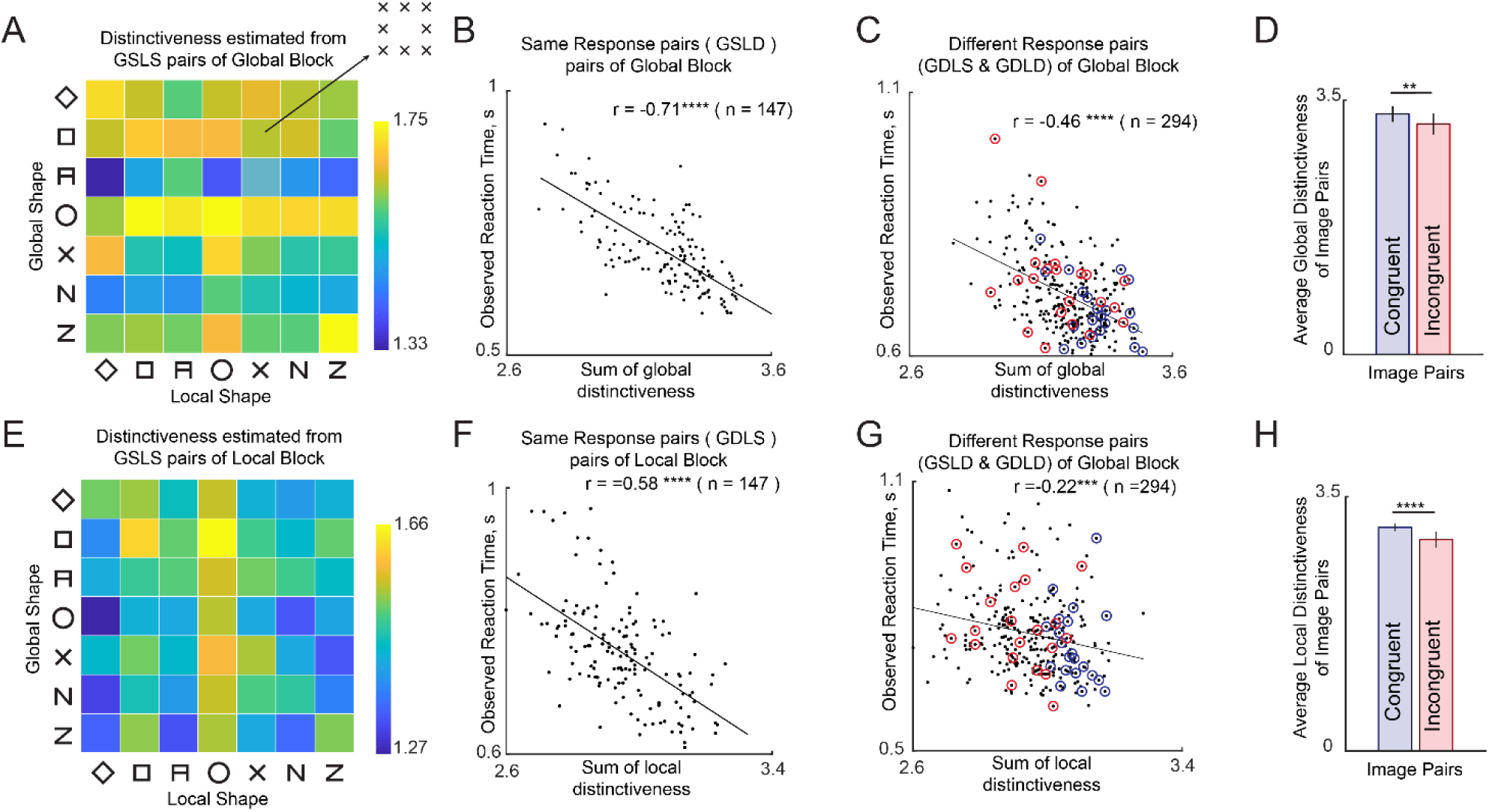
Understanding the contribution of distinctiveness. (A) Global distinctiveness (1/RT) of each hierarchical stimulus, estimated from GSLS pairs in the global block. (B) Observed response times for GSLD pairs in the global block plotted against the net global distinctiveness estimated from panel A. (C) Observed response times for GDLS and GDLD pairs plotted against net global distinctiveness estimated from panel A. (D) Net global distinctiveness calculated for congruent and incongruent image pairs. Error bars represents standard deviation across pairs. (E) Local distinctiveness (1/RT) for each hierarchical stimulus estimated from GSLS pairs in the local block. (F) Observed response times for GDLS pairs in the local block plotted against the net local distinctiveness estimated as in panel D. (G) Observed response times for GSLD & GDLD pairs in the local block plotted against the net local distinctiveness estimated as in panel D. (H) Net local distinctiveness calculated for congruent and incongruent image pairs. Error bar represents standard deviation across pairs.

Having estimated distinctiveness of each image using the GSLS pairs, we asked whether it would predict responses to other pairs. For each image pair containing two different images, we calculated the net distinctiveness as the sum of the distinctiveness of the two individual images. We then plotted the average response times for each GSLD pair (which evoked a “SAME” response) in the global block against the net distinctiveness. This revealed a striking negative correlation (r = −0.71, p < 0.00005; Figure 5B). In other words, subjects responded quickly to distinctive images. We performed a similar analysis for the GDLS and GDLD pairs (which evoke a “DIFFERENT” response). This too revealed a negative correlation (r = −0.46, p < 0.00005 across all GDLS and GDLD pairs, r = −0.38, p < 0.0005 for GDLS pairs; Figure 5C; r = −0.54, p <0.0005 for GDLD pairs). We conclude that image pairs containing distinctive images elicit faster responses.

If distinctiveness measured from GSLS pairs is so effective in predicting responses to all other pairs, we wondered whether it can also explain the incongruence effect. To do so, we compared the net distinctiveness of congruent pairs with that of the incongruent pairs. Indeed, congruent pairs were more distinctive (average distinctiveness, mean ± sd: ± 0.11 s^-1^ for congruent pairs, 3.17 ± 0.14 xx s^-1^ for incongruent pairs, p < 0.005, sign-rank test across 21 image pairs; Figure 5D).

### Effect of distinctiveness on same-different responses in the local block

We observed similar trends in the local block. Again, we estimated distinctiveness for each image as the reciprocal of the response time to the GSLS trials in the local block (Figure 5E). It can be seen that shapes containing a local circle were more distinctive compared to shapes containing a local diamond (Figure 5E). Interestingly, the distinctiveness estimated in the local block was uncorrelated with the distinctiveness estimated in the global block (r = 0.16, p = 0.25).

As with the global block, we obtained a significant negative correlation between the response times for GDLS pairs (which evoked a “SAME” response) and the net distinctiveness (r = −0.58, p < 0.00005; Figure 5F). Likewise, we obtained a significant negative correlation between the response times of GSLD and GDLD pairs (both of which evoke “DIFFERENT” responses in the local block) with net distinctiveness (r = −0.22, p < 0.0005 across 294 GSLD and GDLD pairs; Figure 5G; r = −0.24, p < 0.005 for GSLD pairs; r = −0.18, p < 0.05 for GDLD pairs). We conclude that distinctive images elicit faster responses.

Finally, we asked whether differences in net distinctiveness can explain the difference between congruent and incongruent pairs. As expected, local distinctiveness was significantly larger for congruent compared to incongruent pairs (average distinctiveness, mean ± sd: 3.08 ± 0.05 s^-1^ for congruent pairs, 2.91 ± 0.11 s^-1^ for incongruent pairs, p < 0.00005, sign-rank test across 21 image pairs; Figure 5H).

The above analyses show that distinctiveness directly estimated from response times to identical images can predict responses to other image pairs containing non-identical images. By contrast, there is no direct subset of image pairs that can be used to measure the contribution of image dissimilarity to response times. We therefore devised a quantitative model for the response times to estimate the underlying image dissimilarities and elucidate the contribution of dissimilarity and distinctiveness. Because high dissimilarity can increase response times for “SAME” responses and decrease response times for “DIFFERENT” responses, we devised two separate models for these two types of responses, as detailed below.

### Can “SAME” responses be predicted using distinctiveness and dissimilarity?

Recall that “SAME” responses in the global block are made to image pairs in which the global shape is the same and local shape is different. Let AB denote a hierarchical stimulus made of shape A at the global level and B at the local level. We can denote any image pair eliciting a “SAME” response in global block as AB and AC, since the global shape will be identical. Then according to our model, the response time (SRT) taken to respond to an image pair AB & AC is given by:

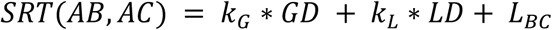

where GD is the sum of the global distinctiveness of AB and AC (estimated from GSLS pairs in the global block), LD is the sum of local distinctiveness of AB and AC (estimated from GSLS pairs in the local block), k_G_, k_L_ are constants that specify the contribution of GD and LD towards the response time, and L_BC_ denotes the dissimilarity between local shapes B and C. Since there are 7 possible local shapes there are only ^7^C_2_ = 21 possible local shape terms. When this equation is written down for each GSLD pair, we get a system of linear equations of the form **y = Xb** where **y** is a 147 x 1 vector containing the GSLD response times, **X** is a 147 x 23 matrix containing the net global distinctiveness and net local distinctiveness as the first two columns, and 0/1 in the other columns corresponding to whether a given local shape pair is present in that image pair or not, and **b** is a 23 x 1 vector of unknowns containing the weights k_G_, k_L_ and the 21 estimated local dissimilarities. Because there are 147 equations and only 22 unknowns, we can estimate the unknown vector **b** using linear regression.

The performance of this model is summarized in Figure 6. The model-predicted response times were strongly correlated with the observed response times for the GSLD pairs in the global block (r = 0.86, p < 0.00005; Figure 6A). These model fits were close to the reliability of the data (rc = 0.84 ± 0.02; see Methods), suggesting that the model explained nearly all the explainable variance in the data. However the model fits do not elucidate which factor contributes more towards response times. To do so, we performed a partial correlation analysis in which we calculated the correlation between observed response times and each factor after regressing out the contributions of the other two factors. For example, to estimate the contribution of global distinctiveness, we calculated the correlation between observed response times and global distinctiveness after regressing out the contribution of local distinctiveness and the estimated local dissimilarity values corresponding to each image pair. This revealed a significant negative correlation (r = −0.81, p < 0.00005; Figure 6A, inset). Likewise, we obtained a significant positive partial correlation between local dissimilarities and observed response times after regressing out the other factors (r = 0.69, p < 0.00005; Figure 6A, inset). However, local distinctiveness showed positive partial correlation (r = 0.30, p = 0.0005) suggesting that locally distinctive shapes slow down responses in the global block. Thus, response times are faster for more globally distinctive image pairs, and slower for more dissimilar image pairs.

**Figure 6.**
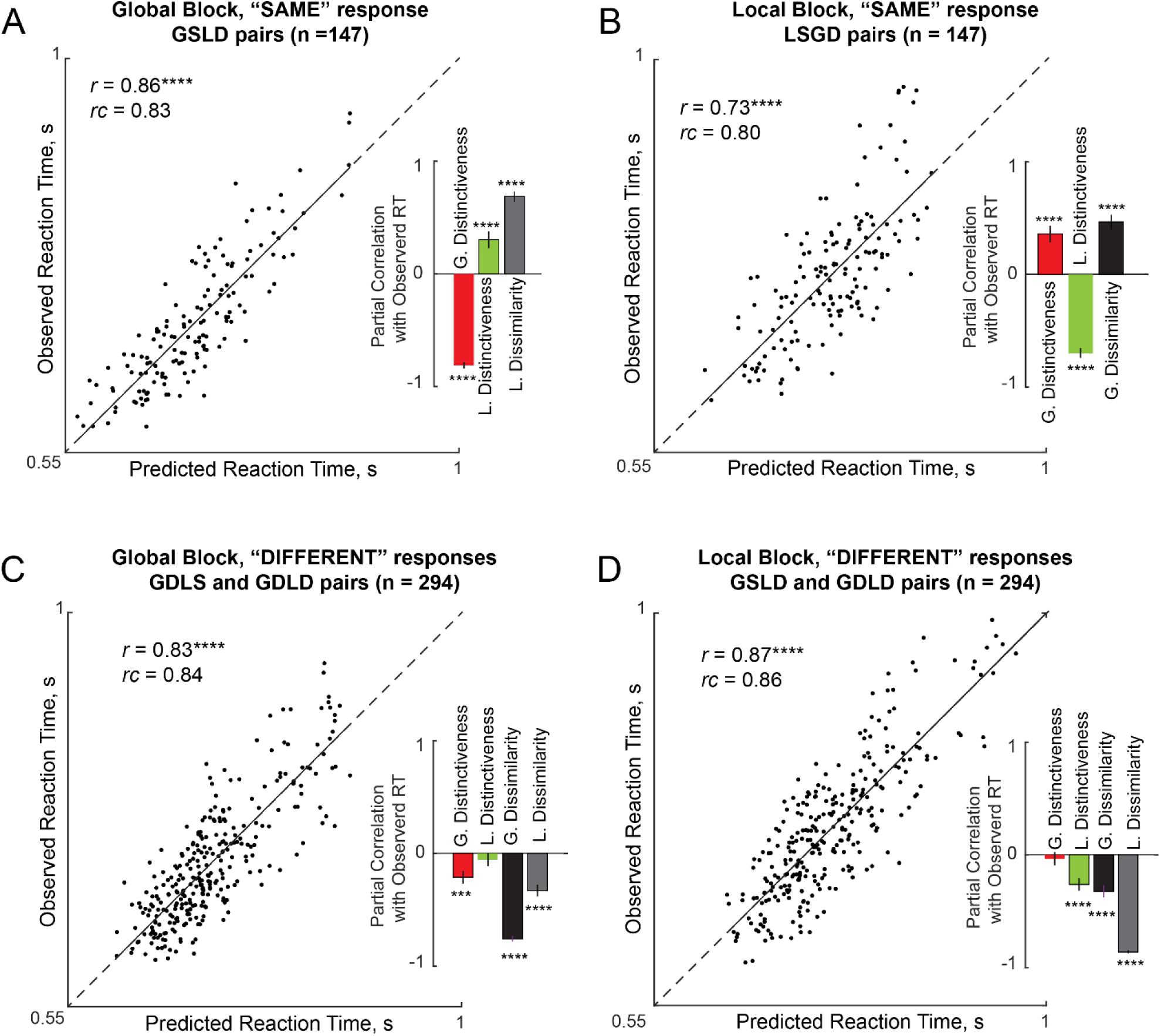
Quantitative model for the Same-Different task. (A) Observed vs predicted response times for “SAME” responses in the global block. *Inset*: partial correlation between observed response times and each factor while regressing out all other factors. Error bars represents 68% confidence intervals, corresponding to ±1 standard deviation from the mean. (B) Same as (A) but for “SAME” responses in the local block. (C) Same as (A) but for “DIFFERENT” responses in the global block. (D) Same as (A) but for “DIFFERENT” responses in the local block.

We obtained similar results for local “SAME” responses. As before, the response time for “SAME” responses in the local block to an image pair (AB, CB) was written as

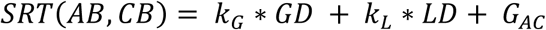

where SRT is the response time, GD and LD are the net global and net local distinctiveness of the images AB and CB respectively, k_G_, k_L_ are unknown constants that specify the contribution of the net global and local distinctiveness and G_AC_ is the dissimilarity between the global shapes A and C. As before this model is applicable to all the LSGD pairs (n = 147), has 23 free parameters and can be solved using straightforward linear regression.

The model fits for local “SAME” responses is depicted in Figure 6B. We obtained a striking correlation between predicted and observed response times (r = 0.72, p < 0.00005; Figure 6B). This correlation was close to the reliability of the data itself (*rc* = 0.80 ± 0.03), suggesting that the model explains nearly all the explainable variance in the response times. To estimate the unique contribution of distinctiveness and dissimilarity, we performed a partial correlation analysis as before. We obtained a significant partial negative correlation between observed response times and local distinctiveness after regressing out global distinctiveness and global dissimilarity (r = −0.70, p < 0.00005; Figure 6B, inset). We also obtained a significant positive partial correlation between observed response times and global dissimilarity after factoring out both distinctiveness terms (r = 0.47, p < 0.00005; Figure 6B, inset). Finally, as before, global distinctiveness showed a positive correlation with local “SAME” responses after accounting for the other factors (r = 0.36, p < 0.00005; Figure 6B inset).

### Can “DIFFERENT” responses be predicted using distinctiveness and dissimilarity?

We used a similar approach to predict “DIFFERENT” responses in the global and local blocks. Specifically, for any image pair AB and CD, the response time according to the model is written as

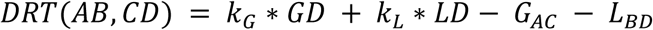

where DRT is the response time for making a “DIFFERENT” response, GD and LD are the net global and net local distinctiveness of the images AB and CD respectively, k_G_, k_L_ are unknown constants that specify their contributions, G_AC_ is the dissimilarity between the global shapes A and C, and LBD is the dissimilarity between the local shapes B and D. Note that, unlike the “SAME” response model, the sign of G_AC_ and L_BD_ is negative because large global or local dissimilarity should speed up “DIFFERENT” responses. The resulting model, which applies to both GDLS and GDLD pairs, consists of 44 free parameters which are the two constants specifying the contribution of the global and local distinctiveness and 21 terms each for the pairwise dissimilarities at the global and local levels respectively. As before, this is a linear model whose free parameters can be estimated using straightforward linear regression.

The model fits for “DIFFERENT” responses in the global block are summarized in Figure 6C. We obtained a striking correlation between observed response times and predicted response times (r = 0.82, p < 0.00005; Figure 6C). This correlation was close to the data reliability itself (rc = 0.84 ± 0.02), implying that the model explained nearly all the explainable variance in the data. To estimate the unique contributions of each term, we performed a partial correlation analysis as before. We obtained a significant negative partial correlation between observed response times and global distinctiveness after regressing out all other factors (r = −0.21, p < 0.0005; Figure 6C, inset). We also obtained a significant negative partial correlation between observed response times and both dissimilarity terms (r = −0.76, p < 0.00005 for global terms; r = −0.33, p < 0.00005 for local terms; Figure 6C, inset). However we note that the contribution of global terms is larger than the contribution of local terms. As before, local distinctiveness did not contribute significantly to “DIFFERENT” responses in the global block (r = −0.06, p = 0.34; Figure 6C, inset). We conclude that “DIFFERENT” responses in the global block are faster for globally distinctive image pairs, and for dissimilar image pairs.

We obtained similar results for “DIFFERENT” responses in the local block for GSLD and GDLD pairs. Model predictions were strongly correlated with observed response times (r = 0.87, p < 0.00005; Figure 6D). This correlation was close to the data reliability (rc = 0.85 ± 0.01) suggesting that the model explained nearly all the variance in the response times. A partial correlation analysis revealed a significant negative partial correlation for all terms except global distinctiveness (correlation between observed RT and each factor after accounting for all others: r = −0.26, p < 0.00005 for local distinctiveness, r = −0.04, p = 0.55 for global distinctiveness, r = −0.32, p < 0.00005 for global terms, r = −0.86, p < 0.00005 for local terms). In contrast to the global block, the contribution of global terms was smaller than that of the local terms. We conclude that “DIFFERENT” responses in the local block are faster for locally distinctive image pairs and for dissimilar image pairs.

### Relation between “SAME” and “DIFFERENT” model parameters

Next we asked whether the dissimilarity terms estimated from “SAME” and “DIFFERENT” responses were related. In the global block, we obtained a significant positive correlation between the local dissimilarity terms (Table 1). Likewise, the global and local terms estimated from “DIFFERENT” responses were significantly correlated (Table 1). In general, only 3 out of 15 (20%) of all possible pairs were negatively correlated, and the median pairwise correlation across all model term pairs was significantly above zero (median correlation: 0.14, p < 0.01). Taken together these positive correlations imply that the dissimilarities driving the “SAME” and “DIFFERENT” responses at both global and local levels are driven by a common underlying shape representation.

**Table 1:**
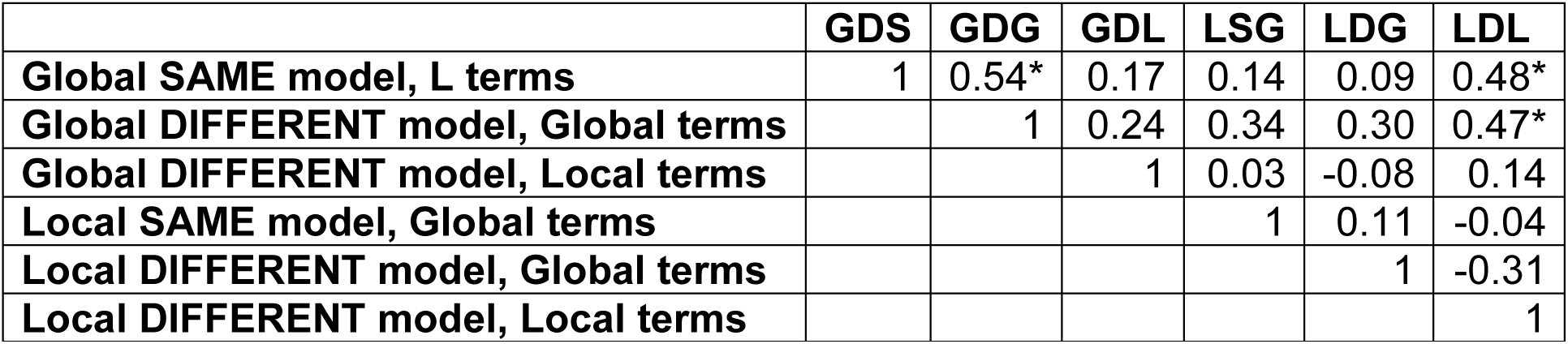
Correlation between estimated dissimilarity terms within and across models. Each entry represents the correlation coefficient between pairs of model terms. Asterisks represent statistical significance (* is p < 0.05). Column labels are identical to row labels but are abbreviated for ease of display.

## EXPERIMENT 2: VISUAL SEARCH

There are two main findings from Experiment 1. First, subjects show a robust global advantage and an incongruence effect in the same-different task. These effects could arise from the underlying categorization process or the underlying visual representation. To distinguish between these possibilities would require a task devoid of categorical judgments. To this end, we devised a visual search task in which subjects have to locate an oddball target among multiple identical distractors, rather than making a categorical shape judgment. Second, responses in the same-different task were explained using two factors: distinctiveness and dissimilarity, but it is not clear how these factors relate to the visual search representation.

We sought to address four fundamental questions. First, are the global advantage and incongruence effects present in visual search? Second, can performance in the same-different task be explained in terms of the responses in the visual search task? Third, can we understand how global and local features combine in visual search? Finally, can the dissimilarity and distinctiveness terms in the same-different model of Experiment 1 be related to some aspect of the visual representations observed during visual search?

### METHODS

#### Subjects

Eight right-handed subjects (6 male, aged 23-30 years) participated in the study. We selected this number of subjects here and in subsequent experiments based on the fact that similar sample sizes have yielded extremely consistent visual search data in our previous studies (Mohan and Arun, 2012; Vighneshvel and Arun, 2013; Pramod and Arun, 2016).

#### Stimuli

We used the same set of 49 stimuli as in Experiment 1, which were created by combining 7 possible shapes at the global level with 7 possible shapes at the local level in all possible combinations.

#### Procedure

Subjects were seated approximately 60 cm from a computer. Each subject performed a baseline motor block, a practice block and then the main visual search block. In the baseline block, on each trial a white circle appeared on either side of the screen and subjects had to indicate the side on which the circle appeared. We included this block so that subjects would become familiar with the key press associated with each side of the screen, and in order to estimate a baseline motor response time for each subject. In the practice block, subjects performed 20 correct trials of visual search involving unrelated objects to become familiarized with the main task.

Each trial of main experiment started with a red fixation cross presented at the centre of the screen for 500 ms. This was followed by a 4 x 4 search array measuring 24° square with a spacing of 2.25° between the centers of adjacent items. Images were were slightly larger in size (1.2x) compared to Experiment 1 to ensure that the local elements were clearly visible. The search array consisted of 15 identical distractors and one oddball target placed at a randomly chosen location in the grid. Subjects were asked to locate the oddball target and respond with a key press (“Z” for left, “M” for right) within 10 seconds, failing which the trial was aborted and repeated later. A red vertical line was presented at the centre of the screen to facilitate left/right judgments.

Search displays corresponding to each possible image pair were presented two times, with either image in a pair as target (with target position on the left in one case and on the right in the other). Thus, there were 49C_2_ = 1,176 unique searches and 2,352 total trials. Trials in which the subject made an error or did not respond within 10 s were repeated randomly later. In practice, these repeated trials were very few in number, because subjects accuracy was extremely high (mean and std accuracy: 98.4% ± 0.7% across subjects).

#### Model fitting

We measured the perceived dissimilarity between every pair of images by taking the reciprocal of the average search time for that pair across subjects and trials. We constructed a quantitative model for this perceived dissimilarity following the part summation model developed in our previous study (Pramod and Arun, 2016). Let each hierarchical stimulus be denoted as AB where A is the shape at the global level and B is the local shape. The net dissimilarity between two hierarchical stimuli AB & CD is given by:

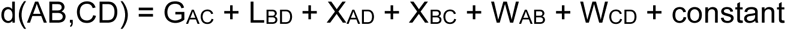

where G_AC_ is the dissimilarity between the global shapes, L_BD_ is the dissimilarity between the local shapes, X_AD_ & X_BC_ are the across-object dissimilarities between the global shape of one stimulus and the local shape of the other, and W_AB_ & W_CD_ are the dissimilarities between global and local shape within each object. Thus there are 4 sets of unknown parameters in the model, corresponding to global terms, local term, across-object terms and within-object terms. Each set contains pairwise dissimilarities between the 7 shapes used to create the stimuli. Note that model terms repeat across image pairs: for instance, the term G_AC_ is present for every image pair in which A is a global shape of one and C is the global shape of the other. Writing this equation for each of the 1,176 image pairs results in a total of 1176 equations corresponding to each image pair, but with only 21 shape pairs x 4 types (global, local, across, within) + 1 = 85 free parameters. The advantage of this model is that it allows each set of model terms to behave independently, thereby allowing potentially different shape representations to emerge for each type through the course of model fitting.

This simultaneous set of equations can be written as **y** = **Xb** where **y** is a 1,176 x 1 vector of observed pairwise dissimilarities between hierarchical stimuli, **X** is a 1,176 x 85 matrix containing 0, 1 or 2 (indicating how many times a part pair of a given type occurred in that image pair) and **b** is a 85 x 1 vector of unknown part-part dissimilarities of each type (corresponding, across and within). We solved this equation using standard linear regression (*regress* function, MATLAB).

The results described in the main text, for ease of exposition, are based on fitting the model to all pairwise dissimilarities, which could result in overfitting. To assess this possibility, we fitted the model each time on 80% of the data and calculated its predictions on the held-out 20%. This too yielded a strong positive correlation across many 80-20 splits (r = 0.85 ± 0.01, p < 0.00005 in all cases), indicating that the model is not overfitting to the data.

## RESULTS

Subjects performed searches corresponding to all possible pairs of hierarchical stimuli (^49^C2 = 1176 pairs). Subjects were highly accurate in the task (mean ± sd accuracy: 98.4% ± 0.7% across subjects).

Note that each image pair in visual search has a one-to-one correspondence with an image pair used in the same-different task. Thus, we have GDLS, GSLD and GDLD pairs in the visual search task. However, there are no GSLS pairs in visual search since these pairs correspond to identical images, and can have no oddball search.

### Is there a global advantage effect in visual search?

We set out to investigate whether there is a global advantage effect in visual search. We compared searches with target differing only in global shape (i.e. GDLS pairs) with equivalent searches in which the target differed only in local shape (i.e. GSLD pairs). Two example searches are depicted in Figure 7A-B. It can be readily seen that finding a target differing in global shape (Figure 7A) is much easier than finding the same shape difference in local shape (Figure 7B).

**Figure 7.**
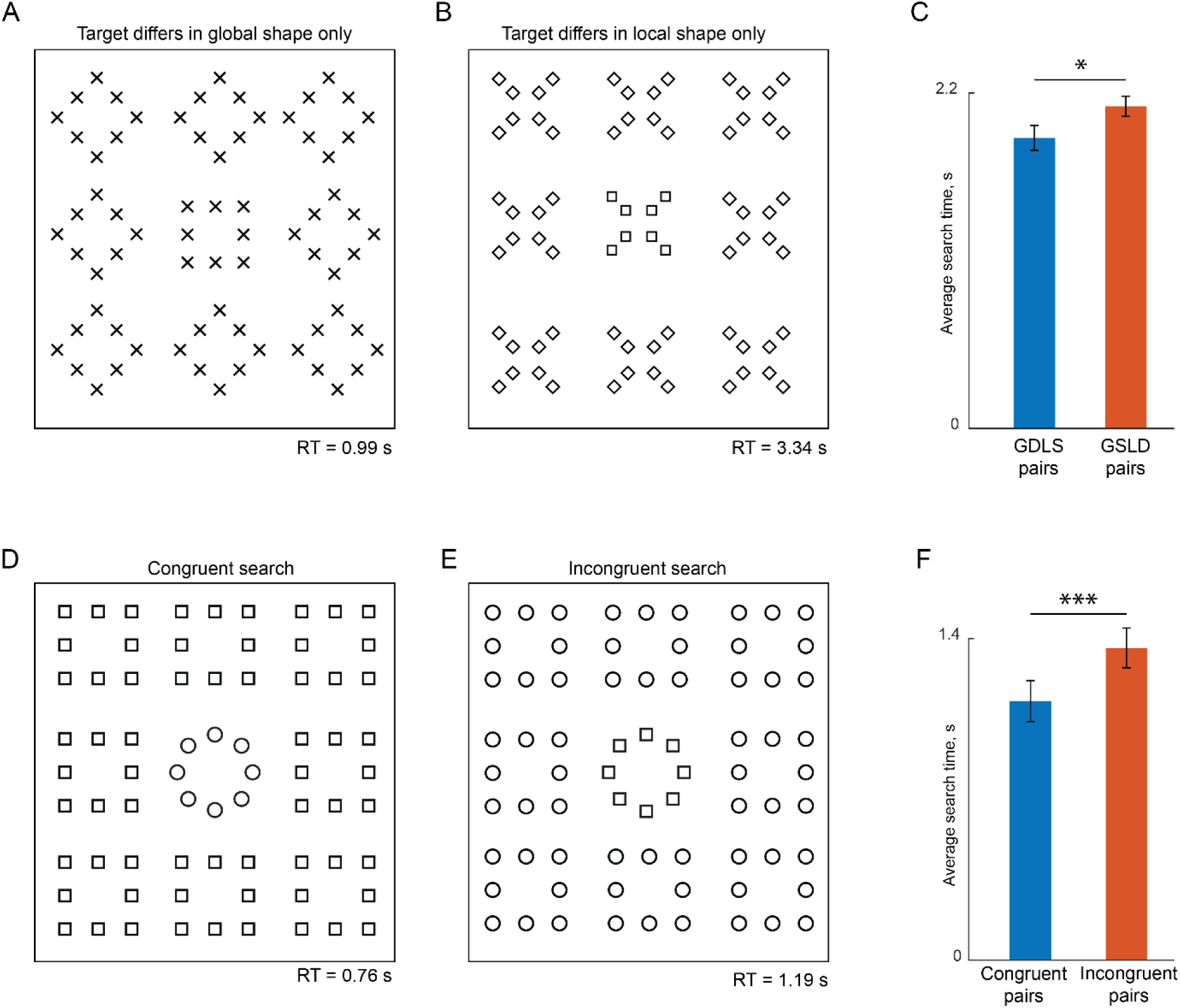
Odd ball visual search task. (A) Example search array with an oddball target differing only in global shape from the distractors. The actual experiment used 4×4 search arrays with stimuli shown as white against a black background. (B) Example search array with an oddball target differing only in local shape from the distractors. (C) Average response times for GDLS and GSLD pairs. Error bars represent s.e.m across subjects. Asterisks indicate statistical significance calculated using a rank-sum test across 147 pairs (* is p < 0.05).. (D) Example search array with two congruent stimuli. (E) Example search array with two incongruent stimuli. (F) Average response time for congruent and incongruent stimulus pairs. Error bars represent s.e.m across subjects. Asterisks indicate statistical significance using an ANOVA on response times (*** is p < 0.0005).

The above observation held true across all GDLS/GSLD searches. Subjects were equally accurate on GDLS searches and GSLD searches (accuracy, mean ± sd: 98% ± 1% for GDLS, 98% ± 1% for GSLD, p = 0.48, sign-rank test across subject-wise accuracy). However they were faster on GDLS searches compared to GSLD searches (search times, mean ± sd: 1.90 ± 0.40 s across 147 GDLS pairs, 2.11 ± 0.56 s across 147 GSLD pairs; Figure 7C).

To assess the statistical significance of this difference, we performed an ANOVA on the search times with subject (8 levels), pairs (7×21 = 147 levels), and hierarchical level (same-global/same-local) as factors. This revealed a significant main effect of hierarchical level (p < 0.00005). We also observed significant main effects of subject and pairs (p < 0.005). All two-way interactions except subject x shape were also significant (p < 0.00005) but these did not alter the general direction of the effect as evidenced by the fact that searches for the same global shape were harder than for the same local shape on average in 82 of 147 pairs (56%) across all subjects. We conclude that searching for a target differing in global shape is easier than searching for a target differing in local shape. Thus, there is a robust global advantage effect in visual search.

### Is there an incongruence effect in visual search?

Next we compared whether searches involving a pair of congruent stimuli were easier than those with incongruent stimuli. Two example searches are shown in Figure 7D-E. It can be readily seen that search involving the congruent stimuli (Figure 7D) is easier than the search involving incongruent stimuli (Figure 7E), even though both searches involve a difference in global shape (circle to square) and a difference in local shape (circle to square).

To establish whether this was true across all 21 searches of this type, we performed an ANOVA on the search times with subject (8 levels), shape pair (^7^C2 = 21 levels) and congruence (2 levels) as factors. This revealed a significant main effect of congruence (average search times: 1.13 s for congruent pairs, 1.36 s for incongruent pairs; p < 0.00005). We also observed a significant main effect of subject and shape pair (p <0.00005), and importantly no significant interaction effects (p > 0.2 for all interactions). We conclude that search involving congruent stimuli are easier than searches involving incongruent stimuli. Thus, there is a robust incongruence effect in visual search.

### Are there systematic variations in responses in the visual search task?

Having established that subjects showed a robust global advantage effect and incongruence effects, we wondered whether there were other systematic variations in their responses as well. Indeed, response times were highly systematic as evidenced by a strong correlation between two halves of the subjects (split-half correlation between RT of odd- and even-numbered subjects: r = 0.83, p < 0.00005).

Previous studies have shown that the reciprocal of search time can be taken as a measure of dissimilarity between the target and distractors. We therefore took the reciprocal of the average search time across all subjects (and trials) for each image pair as a measure of dissimilarity between the two stimuli. Because we performed all pairwise searches between the hierarchical stimuli, it becomes possible to visualize these stimuli in visual search space using multidimensional scaling (MDS). Briefly, multidimensional scaling estimates the 2D coordinates of each stimulus such that distances between these coordinates match best with the observed distances. In two dimensions with 49 hierarchical stimuli, there are only 49 x 2 = 98 unknown coordinates that have to match the ^49^C_2_ = 1,176 observed distances. We emphasize that multidimensional scaling only offers a way to visualize the representation of the hierarchical stimuli at a glance; we did not use the estimated 2D coordinates for any subsequent analysis but rather used the directly observed distances themselves.

The multidimensional scaling plot obtained from the observed visual search data is shown in Figure 8. Two interesting patterns can be seen. First, stimuli with the same global shape clustered together, indicating that these are hard searches. Second, congruent stimuli (i.e. with the same shape at the global and local levels) were further apart compared to incongruent stimuli (with different shapes at the two levels), indicating that searches involving congruent stimuli are easier than incongruent stimuli. These observations concur with the global advantage and incongruence effect described above in visual search.

**Figure 8.**
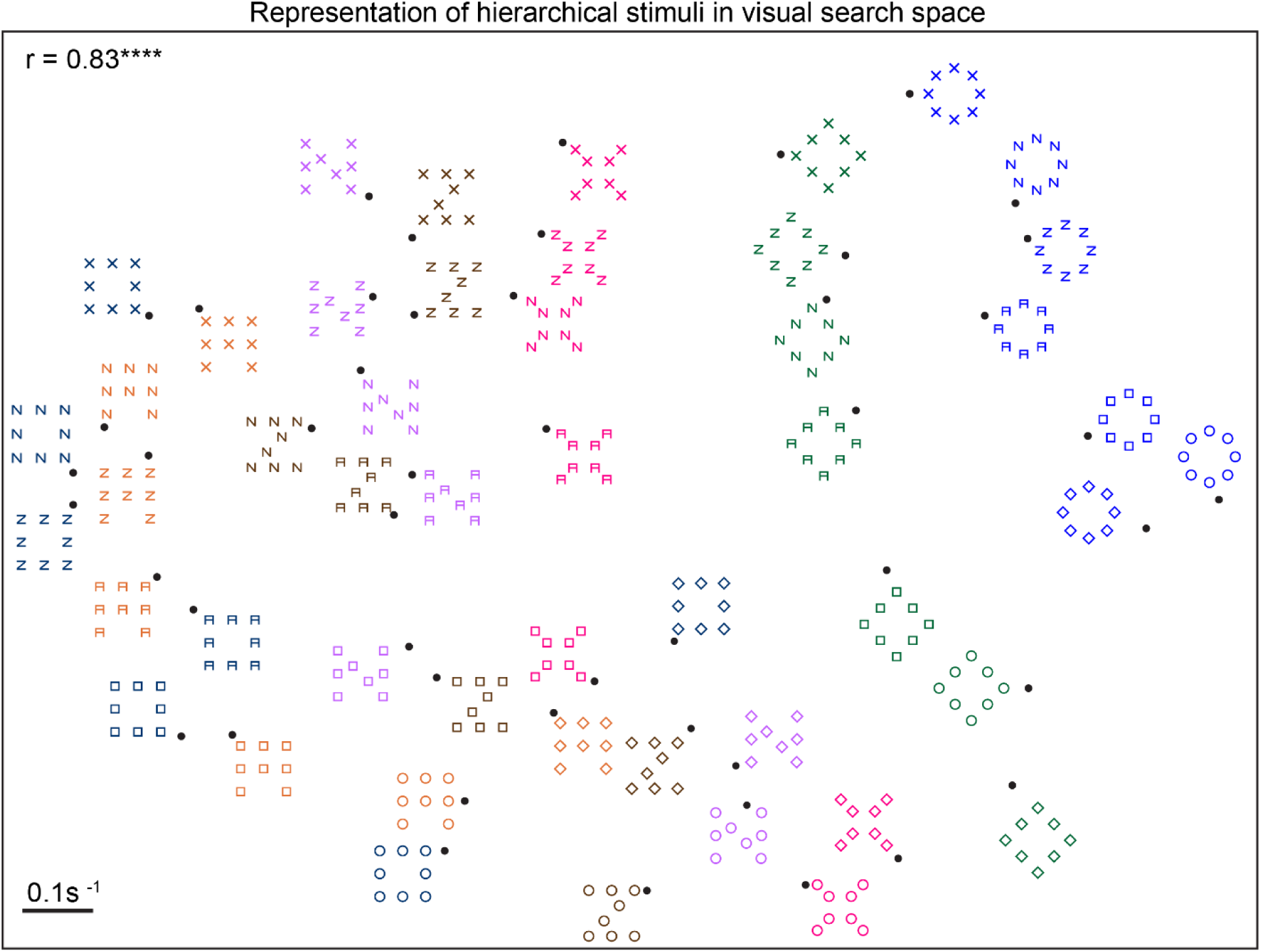
Visualization of hierarchical stimuli in visual search space. Representation of hierarchical stimuli in visual search space, as obtained using multidimensional scaling. Stimuli of the same color correspond to the same global shape for ease of visualization. The actual stimuli were white shapes on a black background in the actual experiment. In this plot, nearby points represent hard searches. The correlation coefficient at the top right indicates the degree of match between the two-dimensional distances depicted here with the observed search dissimilarities in the experiment. Asterisks indicate statistical significance: **** is p < 0.00005.

### How do global and local shape combine in visual search?

So far we have shown that the global advantage and incongruence effects in the same-different task also arise in the visual search task, suggesting that these effects are intrinsic to the underlying representation of these hierarchical stimuli. However, these findings do not provide any fundamental insight into the underlying representation or how it is organized. For instance, why are incongruent shapes more similar than congruent shapes? How do global and local shape combine?

To address these issues, we asked whether search for pairs of hierarchical stimuli can be explained in terms of shape differences and interactions at the global and local levels. To build a quantitative model, we drew upon our previous studies in which the dissimilarity between objects differing in multiple features was found to be accurately explained as a linear sum of part-part dissimilarities (Pramod and Arun, 2014, 2016; Sunder and Arun, 2016). Consider a hierarchical stimulus AB, where A represents the global shape and B is the local shape. Then, according to the model (which we dub the multiscale part sum model), the dissimilarity between two hierarchical stimuli AB & CD can be written as a sum of all possible pairwise dissimilarities between the parts A, B, C and D as follows (Figure 6A):

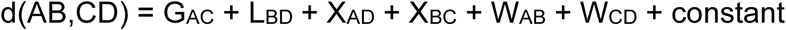

where G_AC_ is the dissimilarity between the global shapes, L_BD_ is the dissimilarity between the local shapes, X_AD_ & X_BC_ are the across-object dissimilarities between the global shape of one stimulus and the local shape of the other, and W_AB_ & W_CD_ are the dissimilarities between global and local shape within each object. Since there are 7 possible global shapes, there are ^7^C_2_ = 21 pairwise global-global dissimilarities corresponding to G_AB_, G_AC_, G_AD_, etc, and likewise for L, X and W terms. Thus in all the model has 21 part-part relations x 4 types + 1 constant = 85 free parameters. Importantly, the multiscale part sum model allows for completely independent shape representations at the global level, local level and even for comparisons across objects and within object. The model works because the same global part dissimilarity G_AC_ can occur in many shapes where the same pair of global shapes A & C are paired with various other local shapes.

### Performance of the part sum model

To summarize, we used a multiscale part sum model that explains the dissimilarity between two hierarchical stimuli as a sum of pairwise shape comparisons across multiple scales. To evaluate model performance, we plotted the observed dissimilarities between hierarchical stimuli against the dissimilarities predicted by the part sum model (Figure 9B). This revealed a striking correlation (r = 0.88, p < 0.00005; Figure 9B). This high degree of fit matches the reliability of the data (mean ± sd reliability: *rc* = 0.84 ± 0.01; see Methods).

**Figure 9.**
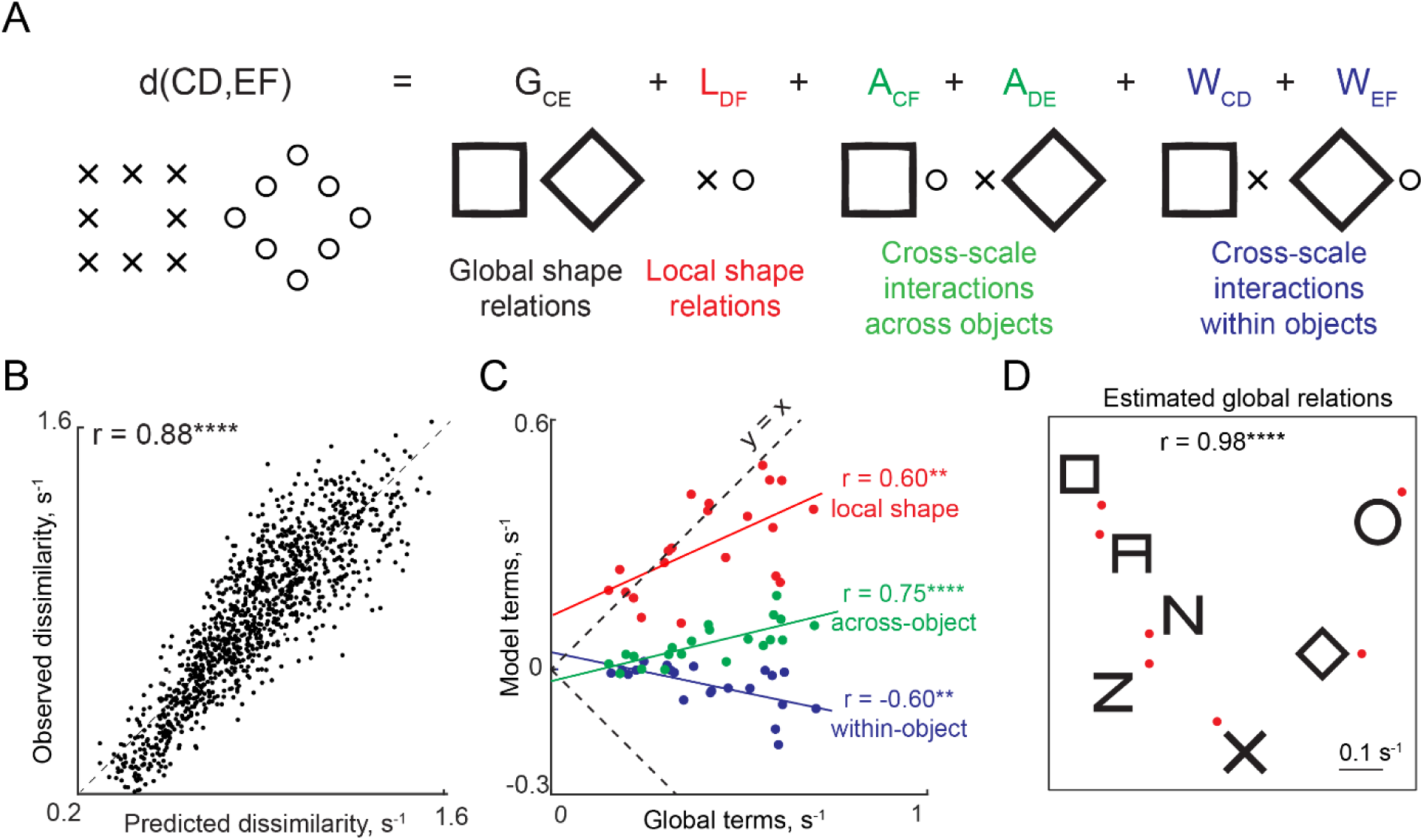
Global and local shape integration in hierarchical stimuli. (A) We investigated how global and local shape combine in visual search using the multiscale part sum model. According to the model, the dissimilarity between two hierarchical stimuli can be explained as a weighted sum of shape differences at the global level, local level and cross-scale differences across and within objects (see text). (B) Observed dissimilarity plotted against predicted dissimilarity for all 1,176 object pairs in the experiment. (C) Local and cross-scale model terms plotted against global terms. Coloured lines indicates the corresponding best fitting line. Asterisks indicate statistical significance: *** is p < 0.0005, **** is p < 0.00005. (D) Visualization of global shape relations recovered by the multiscale model, as obtained using multidimensional scaling analysis.

This model also yielded several insights into the underlying representation. First, because each group of parameters in the part sum model represent pairwise part dissimilarities, we asked whether they all reflect a common underlying shape representation. To this end we plotted the estimated part relations at the local level (L terms), the across-object global-local relations (X terms) and the within-object relations (W terms) against the global part relations (G terms). This revealed a significant correlation for all terms (correlation with global terms: r = 0.60, p < 0.005 for L terms, r = 0.75, p < 0.00005 for X terms, r = −0.60, p < 0.005 for W terms; Figure 9C). This is consistent with the finding that hierarchical stimuli and large/small stimuli are driven by a common representation at the neural level (Sripati and Olson, 2009).

Second, cross-scale within-object (W terms) were negative (average: −0.04, p < 0.005, sign-rank test on 21 within-object terms). In other words, the effect of within-object dissimilarity is to increase overall dissimilarity when global and local shapes are similar to each other and decrease overall dissimilarity when they are dissimilar.

Third, we visualized this common shape representation using multidimensional scaling on the pairwise global coefficients estimated by the model. The resulting plot (Figure 9D) reveals a systematic arrangement whereby similar global shapes are nearby. Ultimately, the multiscale part sum model uses this underlying part representation determines the overall dissimilarity between hierarchical stimuli.

### Can the multiscale model explain the global advantage and incongruence effect?

Having established that the full multiscale part sum model yielded excellent quantitative fits, we asked whether it can explain the global advantage and incongruence effects.

First, the global advantage effect in visual search is the finding that shapes differing in global shape are more dissimilar than shapes differing in local shape. This is explained by the multiscale part sum model by the fact that global part relations are significantly larger in magnitude compared to local terms (average magnitude across 21 pairwise terms: 0.42 ± 0.17 s^-1^ for global, 0.30 ± 0.11 s^-1^ for local, p < 0.005, sign-rank test).

Second, how does the multiscale part sum model explain the incongruence effect? We first confirmed that the model shows the same pattern as the observed data (Figure 10A). To this end we examined how each model term in the model works for congruent and incongruent shapes (Figure 10B). First, note that the terms corresponding to global and local shape relations are identical for both congruent and incongruent stimuli so these cannot explain the incongruence effect. However, congruent and incongruent stimuli differ in the cross-scale interactions both across and within stimuli. For a congruent pair, which have the same shape at the global and local level, the contribution of within-object terms is zero, and the contribution of across-object terms is non-zero, resulting in an overall larger dissimilarity (Figure 10B). In contrast, for an incongruent pair, the within-object terms are negative and across-object terms are zero, leading to a smaller overall dissimilarity.

**Figure 10.**
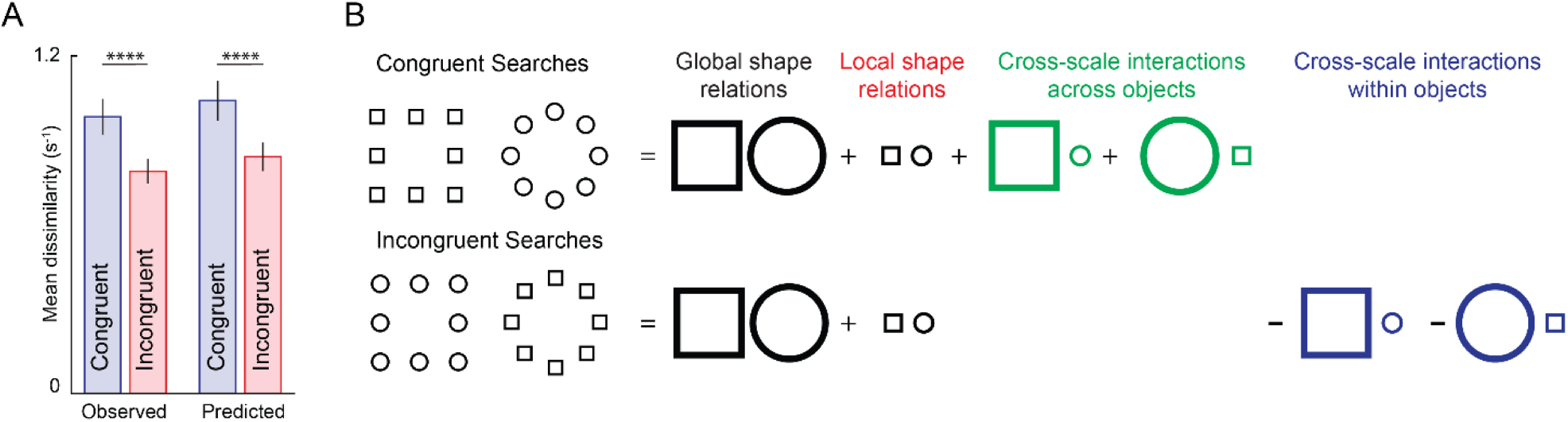
Incongruence effect in visual search. (A) Average dissimilarity for congruent and incongruent image pairs for observed dissimilarities (*left*) and dissimilarities predicted by the multiscale part sum model (*right*). Error bars indicate sd across image pairs. Asterisks indicate statistical significance, as calculated using an ANOVA, with conventions as before. (B) Schematic illustrating how the multiscale model predicts the incongruence effect. For both congruent and incongruent searches, the contribution of global and local terms in the model is identical. However for congruent searches, the net dissimilarity is large because cross-scale across terms are non-zero and within-object terms are zero (since the same shape is present at both scales). In contrast, for incongruent searches, the net dissimilarity is small because across-object terms are zero (since the local shape of one is the global shape of the other) and within-object terms are non-zero and negative.

To summarize, the multiscale model explains qualitative features of visual search such as the global advantage and incongruence effects, and explains visual search for hierarchical stimuli using a linear sum of multiscale part differences. The excellent fits of the model indicate that shape information combines linearly across multiple scales.

### Relating same-different model parameters to visual search

Recall that the responses in the same-different task were explained using two factors, distinctiveness and dissimilarity (Figure 6). We wondered whether these factors are related to any aspect of the visual search representation.

We first asked whether the distinctiveness of each image as estimated from the GSLS pairs in the same-different task is related to the hierarchical stimulus representation in visual search. We accordingly calculated a measure of global distinctiveness in visual search as follows: for each image, we calculated its average dissimilarity (1/RT in visual search) to all other images with the same global shape. Likewise, we calculated local search distinctiveness as the average dissimilarity between a given image and all other images with the same local shape. We then asked how the global and local distinctiveness estimated from the same-different task are related to the global and local search distinctiveness estimated from visual search.

We obtained a striking double-dissociation: global distinctiveness estimated in the same-different task was correlated only with global but not local search distinctiveness (r = 0.55, p < 0.00005 for global search distinctiveness; r = 0.036, p = 0.55 for local search distinctiveness; Figure 11A). Likewise, local distinctiveness estimated in the same-different task was correlated only with local search distinctiveness but not global distinctiveness (r = 0.35, p < 0.05 for local search distinctiveness; r = 0.05, p = 0.76 for global search distinctiveness; Figure 11B).

**Figure 11.**
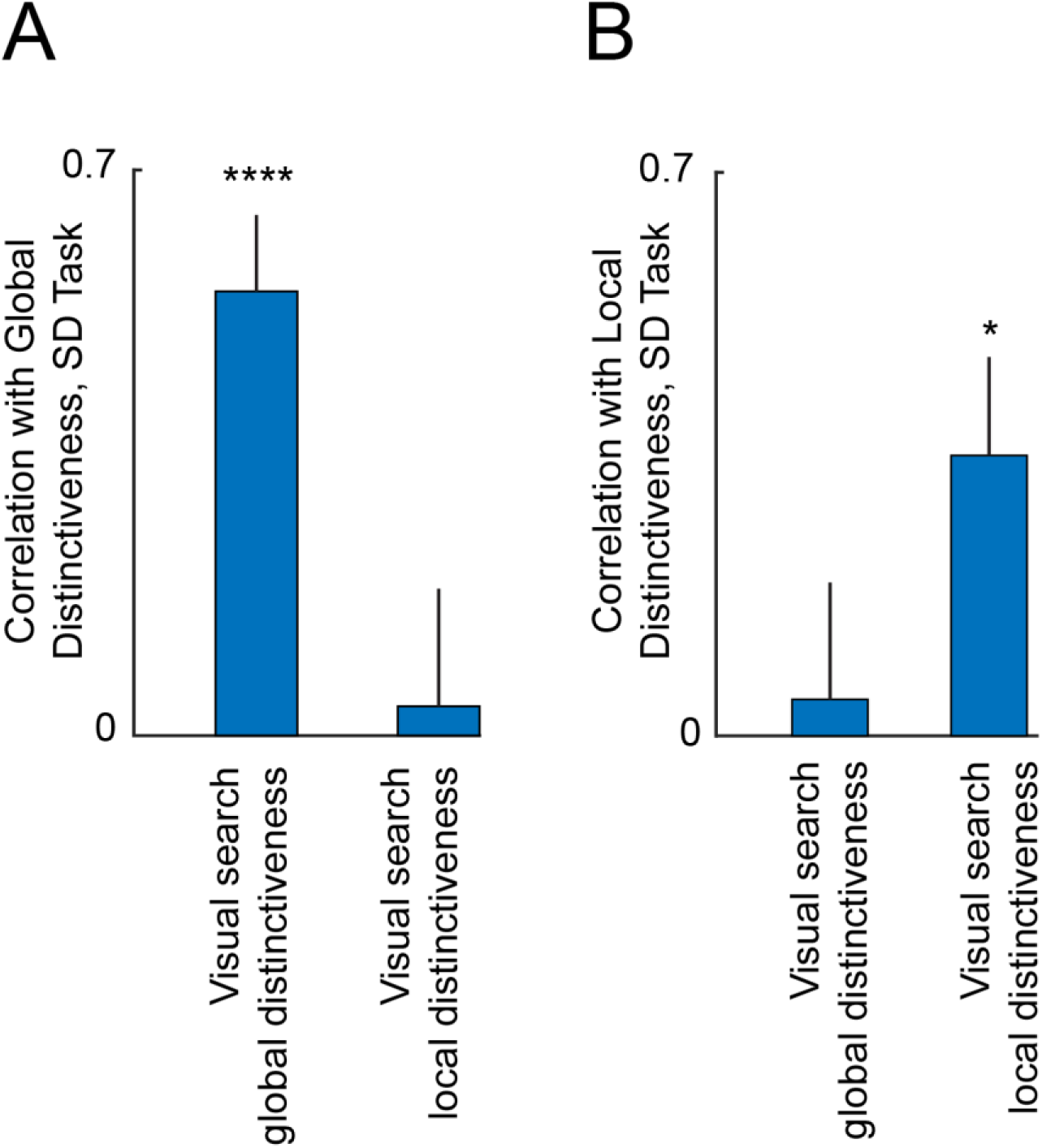
Relation between same-different model parameters and visual search. (A) Correlation between distinctiveness estimated from GSLS trials in the global block of the same-different (SD) task with global and local search distinctiveness. Error bars represents 68% confidence intervals, corresponding to ±1 standard deviation from the mean. (B) Correlation between distinctiveness estimated from GSLS trials in the local block of the same-different task with global and local search distinctiveness.

Next we investigated whether the global and local shape dissimilarity terms estimated from the same-different task were related to the global and local terms in the part-sum model. Many of these correlations were positive and significant (Table 2), suggesting that all dissimilarities are driven by a common shape representation.

**Table 2.** Comparison of model parameters across tasks. Each entry represents the correlation coefficient between model terms estimated from the same-different task and global and local terms from the visual search model. Asterisks represent statistical significance (* is p < 0.05, **** is p < 0.00005 etc).

| Same-Different Model Terms | Correlation with Visual Search Global Terms | Correlation with Visual Search Local Terms |
| --- | --- | --- |
| **Same-Different Task, Global Block** |  |  |
| Same model Local Terms | 0.47* | 0.76**** |
| Different model Global Terms | 0.69**** | 0.82**** |
| Different Model Local Terms | 0.02 | 0 |
| **Same-Different task, Local Block** |  |  |
| Same model Local Terms | 0.37 | 0.11 |
| Different model Global Terms | 0.38 | 0.21 |
| Different Model Local terms | 0.14 | 0.6** |

We conclude that both distinctiveness and dissimilarity terms in the same-different task are systematically related to the underlying representation in visual search.

### Comparison of part-sum model with other models

The above results show that search for hierarchical stimuli is best explained using the reciprocal of search time (1/RT), or search dissimilarity. That models based on 1/RT provides a better account than RT-based models was based on our previous findings (Vighneshvel and Arun, 2013; Pramod and Arun, 2014, 2016; Sunder and Arun, 2016). To reconfirm this finding, we fit RT and 1/RT based models to the data in this experiment. Indeed, 1/RT based models provided a better fit to the data (Section S1).

The above results are also based on a model in which the net dissimilarity is based on part differences at the global and local levels as well as cross-scale differences across and within object. This raises the question of whether simpler models based on a subset of these terms would provide an equivalent fit. However, this was not the case: the full model yielded the best fits despite having more free parameters (Section S1).

### Simplifying hierarchical stimuli

One fundamental issue with hierarchical stimuli is that the global shape is formed using the local shapes, making them inextricably linked. We therefore wondered whether hierarchical stimuli can be systematically related to simpler stimuli in which the global and local shape are independent of each other. We devised a set of “interior-exterior” shapes whose representation in visual search can be systematically linked to that of the hierarchical stimuli, and thereby simplifying their underlying representation. Even here, we found that the dissimilarity between interior-exterior stimuli can be explained as a linear sum of shape relations across multiple scales (Section S2). Moreover, changing the position, size and grouping status of the local elements leads to systematic changes in the model parameters (Section S3-5). These findings provide a deeper understanding of how shape information combines across multiple scales.

## GENERAL DISCUSSION

Classic perceptual phenomena such as the global advantage and incongruence effects have been difficult to understand because they have observed during shape detection tasks, where a complex category judgment is made on a complex feature representation. Here, we have shown that these phenomena are not a consequence of the categorization process but rather are explained by intrinsic properties of the underlying shape representation. Moreover, this underlying representation is governed by a simple rule whereby global and local features combine linearly.

Our findings in support of this conclusion are: (1) Global advantage and incongruence effects are present in a same-different task as well as in a visual search task devoid of any shape categorization; (2) Responses in the same-different task were accurately predicted using two factors: dissimilarity and distinctiveness; (3) Dissimilarities in visual search were explained using a simple linear rule whereby the net dissimilarity is a sum of pairwise multiscale shape dissimilarities. Below we discuss how these results relate to the existing literature.

### Explaining global advantage and incongruence effects

We have shown that the global advantage and incongruence effects also occur in visual search, implying that they are intrinsic properties of the underlying representation. Moreover we show that this representation is organized according to a simple linear rule whereby global and local features combine linearly (Figure 9). This model provides a simple explanation of both effects. The global advantage occurs simply because global part relations are more salient than local relations (Figure 9C). The interference effect occurs because congruent stimuli are more dissimilar (or equivalently, more distinctive) than incongruent stimuli, which in turn is because the within-object part differences are zero for part relations (Figure 10).

Finally, it has long been observed that the global advantage and interference effects vary considerably on the visual angle, eccentricity and shapes of the local elements (Navon, 1977; Navon and Norman, 1983; Kimchi, 1992; Poirel et al., 2008). Our results offer a systematic approach to understand these variations: the multiscale model parameters varied systematically with the position, size and grouping status of the local elements (Section S3-5).

### Understanding same-different task performance

We have found that image-by-image variations in response times in the same-different task can be accurately explained using a quantitative model. To the best of our knowledge, there are no such quantitative models for the same-different task. According to our model, responses in the same-different task are driven by two factors: dissimilarity and distinctiveness.

The first factor is the dissimilarity between two images in a pair. Notably, it has opposite effects on “SAME” and “DIFFERENT” responses. This makes intuitive sense because if images are more dissimilar, it should make “SAME” responses harder and “DIFFERENT” responses easier. It is also consistent with the common models of decision-making (Gold and Shadlen, 2002) and categorization (Ashby and Maddox, 1994; Mohan and Arun, 2012), where responses are triggered when a decision variable exceeds a criterion value. In this case, the decision variable is the dissimilarity.

The second factor is distinctiveness. Response times were faster for images that are more distinctive, i.e. far away from other stimuli. This makes intuitive sense because nearby stimuli can act as distractors and slow down responses. Importantly, the distinctiveness of an image in the global block matched best with its average distance from all other stimuli with the same global shape (Figure 11A). Conversely the distinctiveness in the local block matched best with its average distance from all other shapes with the same local shape (Figure 11B). This finding is concordant with norm-based accounts of object representations (Sigala et al., 2002; Leopold et al., 2006), wherein objects are represented relative to an underlying average. We speculate that this underlying average is biased by the level of attention, making stimuli distinctive at the local or global level depending on the block. Testing these intriguing possibilities will require recording neural responses during global and local processing.

### Linearity in visual search

We have found that the net dissimilarity between hierarchical stimuli can be understood as a linear sum of shape relations across multiple scales. This finding is consistent with our previous studies showing that the net dissimilarity in visual search is a linear sum of elemental feature differences (Pramod and Arun, 2014) as well as of local and configural differences (Pramod and Arun, 2016). Likewise, the net dissimilarity in a search for a target among multiple distractors can be understood as a sum of the dissimilarity of the constituent searches (Vighneshvel and Arun, 2013). More recently, we have demonstrated that knowledge of a forthcoming target adds linearly to bottom-up dissimilarity (Sunder and Arun, 2016). Taken together, these findings suggest that a variety of factors combine in visual search according to a simple linear rule.

## ACKNOWLEDGEMENTS

SPA was supported by Intermediate and Senior Fellowships from the Wellcome-DBT India Alliance (Grant #: 500027/Z/09/Z and IA/S/17/1/503081).

## AUTHOR CONTRIBUTIONS

GJ & SPA designed experiments, GJ collected data, GJ & SPA analysed and interpreted data and wrote the manuscript.

