## Supplementary Text for "How the forest interacts with the trees: Multiscale shape integration explains global and local processing"

**SUPPLEMENTARY MATERIAL**
**for**

**"How the forest interacts with the trees: Multiscale shape integration explains global and local processing" by Georgin Jacob & SP Arun**

**CONTENTS**


SECTION S1. COMPARISON OF RT AND 1/RT MODELS

SECTION S2. SIMPLIFYING HIERARCHICAL STIMULI

SECTION S3. CHANGING ELEMENT SIZE, POSITION & NUMBER

SECTION S4. CHANGING ELEMENT POSITION

SECTION S5. CHANGING ELEMENT GROUPING

### SECTION S1. COMPARISON WITH OTHER MODELS

#### Is search for hierarchical stimuli explained better using RT or 1/RT models?

The results in the main text show that search for hierarchical stimuli is best explained using the reciprocal of search time (1/RT), or search dissimilarity. That models based on 1/RT provides a better account than RT-based models was based on our previous findings (Vighneshvel and Arun, 2013; Pramod and Arun, 2014, 2016; Sunder and Arun, 2016). Here we reconfirmed this finding on the visual search experiment in this study (i.e. Experiment 2).

We tested models based on both search response times (RT) and search dissimilarity (1/RT) to identify the best model that accounts for the data. In each case, we fit the full model, in which the net RT or 1/RT corresponding to the search for two hierarchical stimuli is a weighted sum of shape differences at the global and local level as well as cross-scale terms across and within objects (Figure 7B). Because the two models have the same number of free parameters, we compared their quality of fit directly using their overall correlation with the observed data as well as using their residual errors.

Our main finding is that the 1/RT model outperformed the RT model both in predicting the RT and the 1/RT data in terms of correlations (correlations with 1/RT data: 0.88 and 0.81 for the 1/RT and RT models, $p < 0.00005$, Fisher's z-test; correlations with RT data: 0.88 & 0.87 for the 1/RT and RT models, $p = 0.2$). For a finer-grained comparison between the RT and 1/RT models, we compared their residual errors. Here too, the residual error for the 1/RT model was lower than the RT model for both RT & 1/RT data (average absolute error in RT: 0.21 & 0.28 s for the 1/RT and RT models, $p < 0.00005$, rank-sum test across 1176 observations; average absolute error in 1/RT: 0.09 & 0.13 $s^{-1}$ for the 1/RT and RT models, $p < 0.00005$). We conclude that the 1/RT based model provided a better fit to the search data.

#### Can a simpler multiscale model account for the data?

In the full model described above, the dissimilarity between hierarchical stimuli was taken as a weighted sum of local and global shape differences as well as cross-scale differences both within and across objects. This model yielded excellent fits to the data, but it is possible that a simpler model (using only a subset of these terms) performs just as well.

Comparing the full model with simpler sub-models containing only some types of terms is non-trivial because a complex model will always yield better fits to a given set of data than a simple model by virtue of having more degrees of freedom. Therefore we used a quality of fit measure known as the Akaike's Information Criterion or AICc (Pramod and Arun, 2014, 2016) that penalizes the overall model error by its complexity. The AICc of any model can be calculated as: $AICc = abs\left(N \log\left(\frac{SS}{N}\right) + 2K + \frac{2K(K+1)}{(N-K-1)}\right)$, where N is the number of observations, SS is the sum of squared errors between the model and data across all observations, and K is the number of free parameters in the model. A larger AICc implies a better model.

To compare the quality of fit of two models, we performed a bootstrap analysis. We first resampled the observations with replacement, fit each model and calculated the AICc for each iteration. We then calculated the fraction of bootstrap samples (across 1176 iterations) in which the AICc of one model was larger than that of the other. If this fraction was larger than 95% or smaller than 5% we deemed one model to be superior to the other in terms of the quality of fit.

We fit a number of sub-models that contained various subsets of terms from the full model. Comparing these models on their performance is however not straightforward because some models may have naturally better fits to the data owing to their greater degrees of freedom. We therefore compared the Akaike's Information Criterion or AICc (see above), which takes into account not only the overall residual error between the model predictions and the data, but also penalizes models for having greater degrees of freedom. The results are summarized in Table S1. It can be seen that the full model explains the data better than all sub-models and is superior both in terms of the overall correlation as well as the AICc quality of fit. It can also be seen that global terms contribute the most to the fit, followed by local terms and then by the cross-scale interactions.

| **Model** | ***dof*** | **Model Correlation** | **Quality of fit AICc** *(mean ± sd)* |
|---|---|---|---|
| G | 22 | 0.67**** | 3550 ± 44** |
| L | 22 | 0.45**** | 3114 ± 38** |
| X | 22 | 0.34**** | 2989 ± 46** |
| W | 22 | 0.30**** | 2952 ± 42** |
| GL | 43 | 0.83**** | 4194 ± 53** |
| GX | 43 | 0.71**** | 3619 ± 44** |
| GW | 43 | 0.71**** | 3608 ± 44** |
| LX | 43 | 0.55**** | 3232 ± 47** |
| LW | 43 | 0.52**** | 3175 ± 42** |
| XW | 43 | 0.39**** | 2998 ± 45** |
| GLX | 64 | 0.85* | 4298 ± 52* |
| GLW | 64 | 0.85* | 4291 ± 50* |
| GXW | 64 | 0.74**** | 3676 ± 44** |
| LXW | 64 | 0.59**** | 3250 ± 49** |
| **Full Model (GLXW)** | **85** | **0.88** | **4430 ± 52** |

**Table S1. Comparison of submodels with the full 1/RT model.** In each case the 1/RT model containing a subset of the model terms was fit to the full set of 1176 search dissimilarities. The best model, depicted in **bold face**, was the full model containing global (G), local (L), cross-scale across object (X) and cross-scale within object (W) terms. Asterisks in the model correlation column indicate the statistical significance of comparing each model with the best model using a Fisher's z-test on correlation coefficients (* is $p < 0.05$, ** is $p < 0.005$ etc). Asterisks in the AICc column indicate statistical significance of comparing each model with the best model, calculated as the fraction of bootstrap samples in which the AICc was larger than the AICc of the best model.

### SECTION S2: SIMPLIFYING HIERARCHICAL STIMULI

Having characterized how global and local shape combine in hierarchical stimuli, we wondered whether we can obtain further insights by varying their component properties. One fundamental issue with hierarchical stimuli is that the global shape is formed using the local shapes, making them inextricably linked. We therefore wondered whether hierarchical stimuli can be systematically related to simpler stimuli in which the global and local shape are independent of each other.

These simpler stimuli are shown in Figure S1A. For each hierarchical stimulus, we created an equivalent "interior-exterior" stimulus in which an external contour with the same shape as the global shape encloses a random arrangement of interior elements with the same number and local shape (Figure S1A). We repeated this for two element sizes because the grouping of local elements into a global shape is affected by size (Figure S1B). This design allowed us to ask whether feature integration is similar in hierarchical stimuli compared to the interior-exterior stimuli.

#### METHODS

*Subjects.* Eight right-handed human subjects (7 male, aged 23-28 years) participated in the study. All other details were as in Experiment 2.

*Stimuli.* We designed hierarchical stimuli (H) and matched interior-exterior (IE) stimuli. The hierarchical stimuli were created by combining 5 shapes at the local and global levels in all possible combinations, resulting in a total of 25 stimuli. The interior-exterior (IE) stimuli were derived from the hierarchical stimuli by arranging the local shapes in a fixed configuration, and replacing the global form of the hierarchical stimulus by a solid closed contour (Figure S1A). Shapes were chosen such that their exterior/global version was large enough to accommodate 8 local shapes without intersecting with the local shapes. To investigate how the size of the local elements influences the overall dissimilarity, we created a new set of hierarchical and interior-exterior stimuli in which the local elements were 75% of their original size (size 2; Figure S1B). Thus there were four sets of 25 stimuli used in the experiment (hierarchical and interior-exterior at 2 sizes each). Subjects performed visual search involving all possible pairs of stimuli within each set, with the result that there were ${}^{25}C_2 \times 4 = 1200$ unique searches in the experiment. All other details were identical to Experiment 2.

*Model fitting.* We fit the multiscale model to all 300 searches corresponding to each stimulus set. Since there were 5 unique parts, there were ${}^{5}C_2 = 10$ model parameters for each group of global, local, across and within terms. Together with a constant term, the multiscale model consisted of 41 free parameters in all. All other details are as in Experiment 2.

#### RESULTS

Subjects performed visual search using matched hierarchical stimuli and interior-exterior stimuli at two local element sizes (Figure S1A-B). For ease of exposition, we first describe results for the hierarchical and interior-exterior stimuli at the larger size (size 1), and then describe the effect of changing local element size (size 1 vs 2).

**Is the representation of interior-exterior stimuli similar to hierarchical stimuli?**

Subjects were highly consistent in their performance on searches involving both hierarchical and interior-exterior stimuli (split-half correlation between RT for odd and even subjects: 0.79 & 0.93 for hierarchical and interior-exterior stimuli both of size 1, $p < 0.00005$).

To visualize the underlying representation, we performed a multidimensional scaling analysis on the average search dissimilarities for each set separately. As before, the hierarchical stimuli tended to group according to the global shape (Figure S1C). This trend was even more evident for the interior-exterior stimuli (Figure S1D). Thus, hierarchical and interior-exterior stimuli have qualitatively similar representations.

To quantify these observations, we directly compared the pairwise dissimilarities between hierarchical stimulus pairs and interior-exterior pairs. This revealed a significant positive correlation ($r = 0.65$, $p < 0.0005$; Figure S1E). We note that this correlation is only modest even though the interior-exterior dissimilarities and hierarchical dissimilarities were themselves highly consistent. This implies that there are subtle representational differences between the two sets. We therefore wondered whether the multiscale model would be able to account for these differences. This was indeed the case: multiscale model predictions on both sets were excellent ($r = 0.91$ & $0.96$ for hierarchical and interior-exterior sets; Figure S1F). These correlations were virtually the same as the reliability of the data itself ($rc = 0.89 \pm 0.008$ for hierarchical stimuli; $rc = 0.90 \pm 0.003$ for interior-exterior stimuli). Thus, the multiscale model explains nearly all the explainable variance in the data. This in turn implies that whatever subtle representational differences exist between hierarchical and interior-exterior stimuli must arise from systematic differences in their model parameters.

We therefore compared the model parameters for the two sets – and observed several interesting patterns (Figure S1G). First, model terms corresponding to global shape differences were stronger in the interior-exterior stimuli (average magnitude: 0.47 & 0.94 for hierarchical and interior-exterior, $p < 0.005$, sign-rank test on 10 global terms). This is as expected given the stronger clustering by global shape for the interior-exterior stimuli. However the global terms for hierarchical and interior-exterior stimuli were significantly correlated, indicating that the underlying representation is similar ($r = 0.73$, $p = 0.016$). This correlation was even higher across all model terms ($r = 0.85$, $p < 0.00005$ across 41 model terms for hierarchical and interior-exterior stimuli).

Second, model parameters corresponding to local shape differences and cross-scale interactions were weaker in the interior-exterior stimuli (average magnitude of local terms: 0.13 & 0.045 for hierarchical and interior-exterior, $p < 0.005$, sign-rank test; cross-scale across object terms: 0.1 & 0.03, $p < 0.005$; cross-scale within-object: 0.18 & 0.06, $p < 0.005$; Figure S1G). Third, as before, in both sets, model parameters corresponding to local and cross-scale terms were generally correlated with the global terms in the same way as in Experiment 1 (correlation with global terms for hierarchical stimuli across ${}^5C_2 = 10$ shape pairs: $r = 0.9$, $p < 0.005$ for local, $r = 0.86$, $p < 0.005$ for across and $r = -0.57$ , $p = 0.08$ for within terms; for interior-exterior stimuli: $r = 0.75$, $p < 0.05$ for local, $r = 0.33$, $p = 0.33$ for across and $r = -0.19$, $p = 0.5$ for within terms). These correlations indicate that model parameters are driven by a common shape representation.

**How do model parameters change when local shapes are made smaller?**

Next we asked how the estimated model parameters of the hierarchical and interior-exterior stimuli change if the local elements became smaller. In general search times with larger local elements was faster (average search times: 1.72 and 1.90 s for hierarchical stimuli size 1 and 2, $p < 0.005$, sign-rank test on mean response times; 1.32 and 1.37 for interior-exterior stimuli size 1 & 2, $p = 0.83$, sign-rank test). The multiscale model again yielded excellent fits at this size too (correlation between observed & predicted dissimilarity for size 2: $r = 0.93$ for hierarchical stimuli, $r = 0.96$ for interior-exterior stimuli, $p < 0.00005$ in both cases). Importantly, model parameters changed systematically when local elements were smaller (Figure S1H). These changes were similar for both sets of stimuli, suggesting that local element size influences both stimuli similarly. The general pattern is that, when local elements decrease in size, global terms become larger whereas local and cross-scale terms become weaker (Figure S1H).

To summarize, the multiscale model provided excellent fits for searches involving both hierarchical and interior-exterior stimuli even across changes in local element size. Both stimuli were driven by a common underlying shape representation, and their differences were explained by systematic differences in model parameters. The differences in the model parameters indicate that interior-exterior stimuli have more salient exterior shapes with weaker local and cross-scale interactions. The weaker local and cross-scale interactions could be due to the increased salience of the global shape or due to the greater proximity of the local shapes to each other. In subsequent experiments we designed stimuli to distinguish between these possibilities.

**How do model parameters change with local shape properties?**

The above findings suggest that the multiscale part sum model parameters change systematically with local element size. To further investigate how model parameters change with other local shape properties, we varied the size, position, number and grouping status of the local elements in the interior-exterior stimuli (Sections S2-4). We obtained excellent model fits in all cases, and model parameters varied systematically with these manipulations.

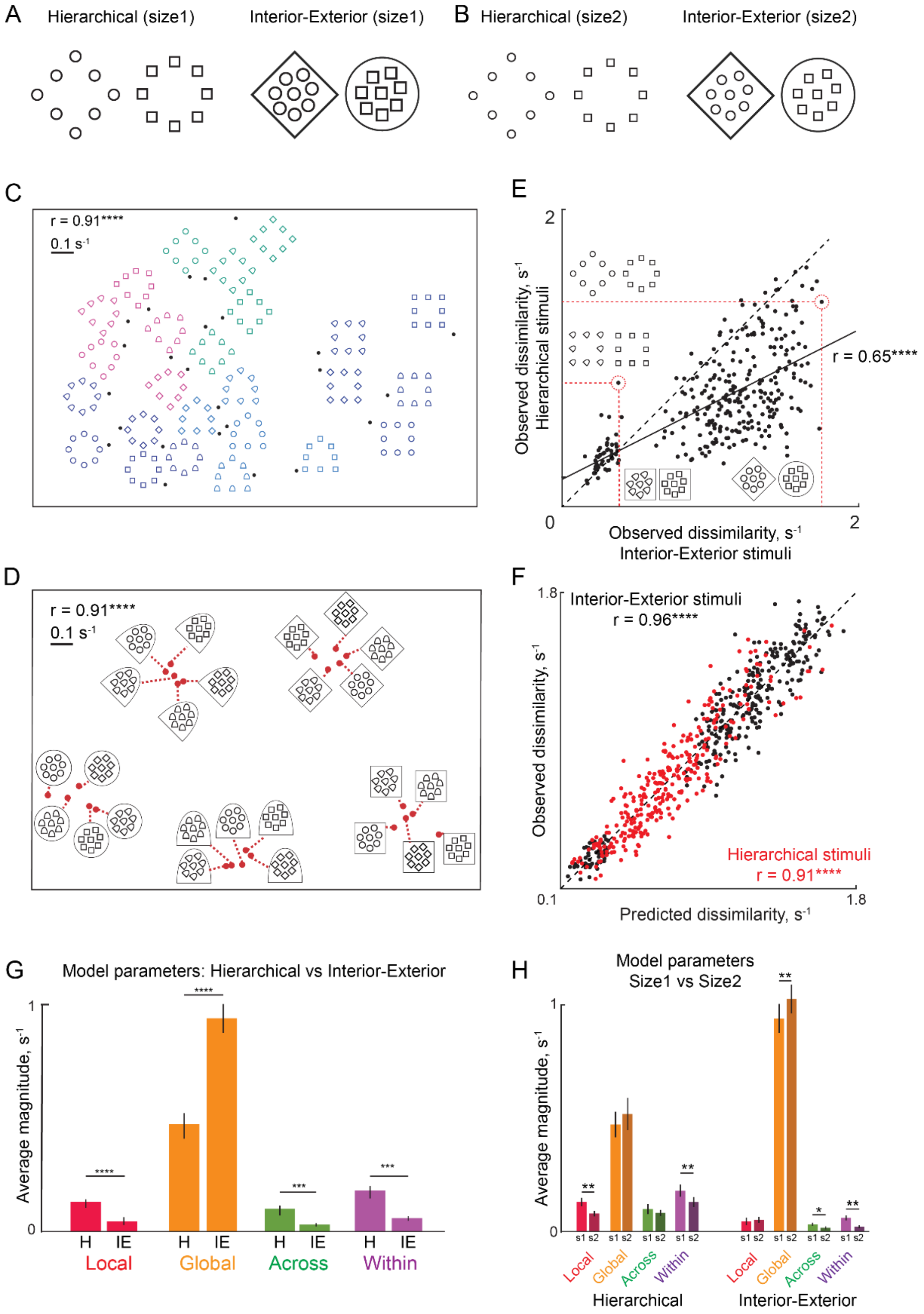


**Figure S1. Simplifying hierarchical stimuli into interior-exterior stimuli**

(A) Example pair of hierarchical stimuli and interior-exterior stimuli. Understanding global and local shape integration in hierarchical stimuli is complicated by the fact that the global shape is inextricably linked to and formed by the local shape. We attempted to simplify each hierarchical stimulus into an “interior-exterior” stimulus in which the external contour matches the global shape and the shape and number of the internal elements matches the local shape.

(B) Example hierarchical and interior-exterior stimulus pairs with smaller size elements (size 2).
(C) Visualization of the underlying shape representation for hierarchical stimuli (Size 1), as obtained using multidimensional scaling.
(D) Same as (C) but for the matched interior-exterior stimuli (Size 1).
(E) Observed dissimilarities for hierarchical pairs plotted against that of Interior-Exterior pairs (Size 1), with example pairs highlighted (*red dotted lines*). The solid line represents the best-fitting straight line and the dotted line is the y = x line.
(F) Observed versus predicted dissimilarity for hierarchical stimuli (*red*) and interior-exterior stimuli (*black*) for Size 1.
(G) Average magnitude of model terms for hierarchical stimuli (H) and Interior-Exterior (IE) stimuli for Size 1. Note that within-object terms are generally negative but their magnitude is depicted for ease of comparison. Asterisks indicate statistical significance as calculated using a sign-rank test on the model parameters, with conventions as before.
(H) Average magnitude of model parameters for Size 1 vs Size 2 for both hierarchical (H) and interior-exterior (IE) stimuli. Asterisks indicate statistical significance as calculated using a sign-rank test on the model parameters, with conventions as before.

### SECTION S3. CHANGING ELEMENT SIZE, POSITION & NUMBER

In Experiment 3, we demonstrated that hierarchical and interior-exterior stimuli are driven by a common shape representation. Here we manipulated the position, size and numerosity of local elements in highly simplified interior-exterior stimuli to understand how these changes affect the overall representation.

#### METHODS

*Subjects.* Eight right-handed human subjects (5 male, aged 21-28 years) participated in the study. All other details were as in Experiment 1.

*Stimuli.* We created four sets each containing 25 stimuli. Set 1 was a reference set containing a single exterior shape and a single interior shape (Figure S2). In Set 2, all stimuli were identical to Set 1 except that the interior shape was shifted to the left. In Set 3, the interior shape was double the size of the Set 1 stimuli. In Set 4, there were two local elements of the same size as in Set 1.

*Procedure.* Subjects performed searches involving all pairwise stimuli within each set. Thus in all there were $^{25}C2$ x 4 sets = 1200 searches. Subjects performed 98.3 ±0.001% correct trials for each unique search. All details were identical to Experiment 1 except that the Set 1 stimuli measured 3.4° along the longest dimension, and the inter-item spacing was slightly smaller at 3.35°.

#### RESULTS

We measured visual search performance on four sets of stimuli in which local elements were varied in position, size and number (Figure S2A). Subjects were highly consistent in their search performance across all four sets (split-half correlation between RT of odd- and even-numbered subjects: r = 0.92, 0.92, 0.87 & 0.89 for Sets 1-4 respectively, $p < 0.00005$ in all cases). Observed dissimilarity was also highly correlated across sets, indicating that the underlying shape representations are very similar (Figure S2B).

To visualize the underlying shape representation we performed a multidimensional scaling as before. The resulting plot for Set 1 is shown in Figure S2C. It can be seen that stimuli with the same global shape cluster together, indicating that these are hard searches.

As before, the multiscale model yielded excellent fits to the data (r = 0.95, 0.95, 0.93 & 0.94 for Sets 1-4 respectively, $p < 0.00005$ in all cases; Figure S2D), implying that variations in the underlying representation across sets due to local element properties are captured by systematic changes in model parameters. These changes are summarized in Figure S2E. The most obvious pattern is that the global terms are substantially larger than all other model terms, indicating that search difficulty is dominated by differences in global shape (Figure S2E). However model parameters varied systematically across the four sets, as discussed below.

We first asked what happens to the shape representation with a change in the local element position (Set 1 vs Set 2). Interestingly, when the local element is shifted away from the centre, local terms became smaller and within-object interactions increased (Figure S2E). However, the underlying shape representation was extremely similar, as evidenced by a strong correlation between the global terms across both sets (r = 0.96, $p < 0.00005$).

Next, we analysed how the shape representation changes when the local elements increase in size (Set 1 vs Set 3). We found that global terms decreased, whereas cross-scale interactions both across and within objects increased (Figure S2E). Because some of these changes tend to increase dissimilarity whereas others will cause a decrease, the overall dissimilarity is unlikely to change. Indeed, search times were not systematically different across the two sets (average search times: 0.88 & 0.90 s for Sets 1 & 3 respectively, $p = 0.11$, rank sum test across 300 searches). Thus, increasing local element size increases cross-scale interactions across hierarchical levels.

We then asked how the shape representation changes when the number of local elements increases from 1 to 2. The only significant change was that cross-scale across-object terms were larger in Set 4 compared to Set 1 (Figure S2E). Set 3 is also an interesting comparison with Set 4 because the total area of the local elements is the same in both Sets. Here, we found that global terms were larger, but within-object interactions were smaller for two local elements (Set 4) compared to one large element (Set 3). Taken together, these changes mean that increasing the number of elements increases cross-scale interactions compared to a single small element, but the net increase is still much smaller compared to having a single large element of the same size.

To summarize, visual search for interior-exterior stimuli across changes in local element position, size and number is explained extremely well by the multiscale model. Moving local elements away from the centre (closer to the exterior shape), increasing their number, or increasing size all led to increased cross-scale interactions.

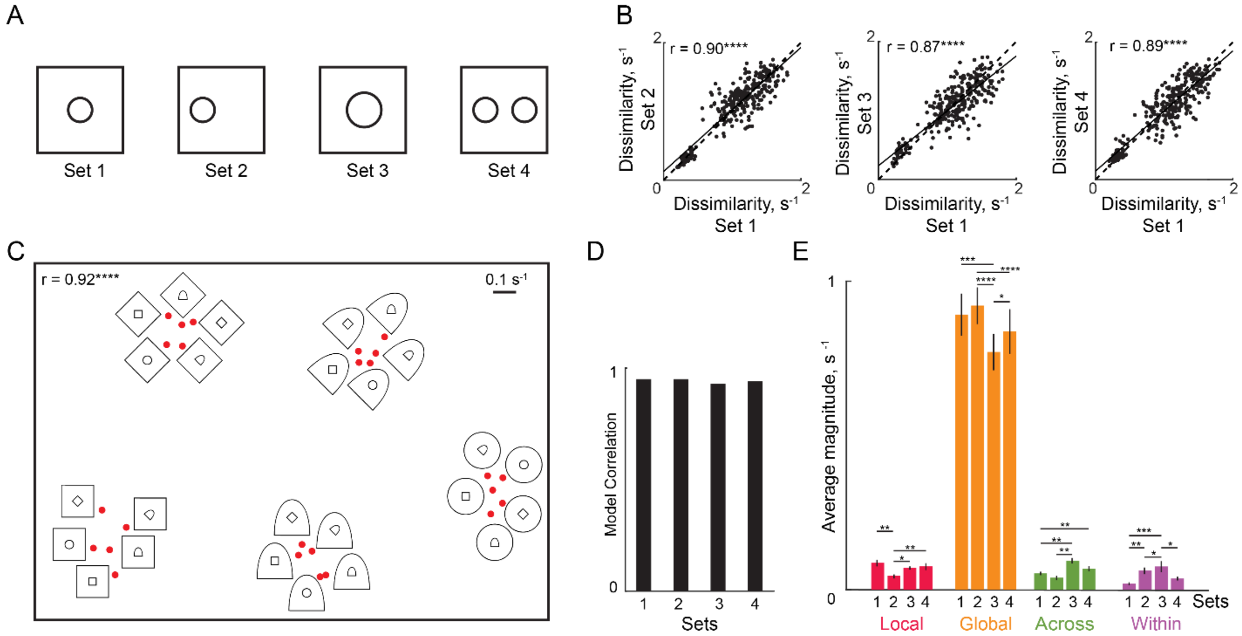


**Figure S2: Effect of local element position, size and number**

(A) Example stimuli from Sets 1-4. Set 1 is the reference, with the local shape at the centre of the exterior contour. In Set 2 the interior shape is shifted away from the centre. In Set 3, the local shape is doubled in size. In Set 4, two local shapes are placed equidistant from the centre on either side.

(B-D) Observed dissimilarity of all 300 pairs of stimuli in each set plotted against Set 1. The *solid line* is the best-fitting line and the *dotted line* represents the unit line (y = x).

(E) Visualization of the underlying shape representation for the reference set (Set 1), as obtained using multidimensional scaling. All conventions are as before.

(F) Correlation between predicted and observed dissimilarities for each set.

(G) Average magnitude of model parameters for Sets 1-4. Asterisks represent statistical significance assessed using a signed-rank test: * is $p < 0.05$, ** is $p < 0.005$. All comparisons are not significant ($p > 0.05$) unless marked with an asterisk.

### SECTION S4: CHANGING ELEMENT POSITION

In Section S2, we showed that moving a local element away from the center of an exterior shape tended to increase cross-scale interactions. Here, we explored this issue further by asking what would happen if the local element was moved even further to intersect the exterior shape or even be located outside it.

#### METHODS

*Subjects.* Eight right-handed human subjects (6 male, aged 20-26 years) participated in the study. All other details are as in Experiment 1.

*Stimuli.* We created interior-exterior stimuli with a single local shape whose bounding box was quarter the area of that of the global shape (FIgure S3A). We also modified the local shapes to be larger in size compared to the Experiment 3 so as to increase the salience of the local and cross-scale terms and reduce the dominance of the global terms. We created four sets of 25 stimuli each by combining five shapes in the interior with the same five shapes for the exterior contour in all possible ways. The four sets were identical except for the position of the local shape: it could be at the centre (Set 1), between the centre and the left edge (Set 2), centred on the global shape contour (Set 3) and finally located outside the exterior shape (Set 4). These are depicted in Figure S3A. To avoid novel conjunctions with the exterior shapes as the interior shape is moved, we used shapes with vertical edges on both left and right sides, with the result that all shapes differed only in contours on the top or bottom sides. The interior shape was presented in a green colour to facilitate grouping particularly for Set 3 where the interior and exterior contours overlap.

*Procedure.* Subjects performed an oddball search task on 4 x 4 search arrays as before, with the largest item measuring 4.1°. All other details are same as that of Experiment 2.

#### RESULTS

In this experiment, subjects performed searches on sets of interior-exterior stimuli in which the center element varied in position. Subjects were highly consistent in their search performance on stimuli in each set (split-half correlation between RT of odd- and even-numbered subjects: $r = 0.82$, 0.87, 0.80 & 0.81 for Sets 1-4 respectively, $p < 0.0005$ in all cases). The observed dissimilarity was also extremely similar across Sets, suggesting that the underlying shape representation is qualitatively similar (Figure S3B). However, search difficulty varied systematically across sets (average search times: 2.00, 1.97, 2.35 and 2.13 s for Sets 1-4), with Set 2 being the easiest ($p < 0.05$, rank-sum test across 300 searches of Set 2 with all other sets) and Set 3 being the hardest ($p < 0.00005$, rank-sum test across 300 searches of Set 3 with all other sets).

To visualize the underlying shape representation, we performed multidimensional scaling as before. In the resulting plot, shown for Set 1 (Figure S3C), it can be seen that stimuli are still clustered according to their global shapes but the grouping is not as strong as in Experiment 3 (i.e. compared to Figure S2C).

As before, the multiscale model yielded excellent fits to the data ($r = 0.93$, 0.93, 0.93 & 0.92 for Sets 1-4 respectively, $p < 0.00005$ in all cases; FIgure S3D), implying that variations in the shape representation due to local element position is captured by systematic changes in model parameters. These changes are summarized in FIgure S3E. Unlike the previous experiment (Section S3) where global terms dominated all others, global terms were comparable in magnitude to other model terms and did not vary with element position (FIgure S3E). We observed systematic changes in model parameters across sets, as detailed below.

We observed a non-monotonic change in model parameters across sets: local terms became larger from Set 1 to Set 2 as in the previous experiment but became much smaller for Set 3 (where the local & global contours overlap) and increased again from Set 3 to 4 (Figure S3E). Cross-scale across-object terms also followed the same pattern although they did not show as big a drop for Set 3 as the local terms (Figure S3E). Cross-scale within-object interactions were strongest when the local shape was at the centre and decreased in magnitude as its position shifted to the left.

Sets 2 & 4 are an interesting comparison because the local shape is equally far away from the edge of the exterior contour, but different both in terms of being inside vs outside as well as distance from the centre of the exterior contour. Compared to Set 2, local and cross-scale terms (both across and within) were smaller in Set 4 (Figure S3E).

To summarize, visual search for interior-exterior stimuli is explained extremely well by the multiscale model across changes in local element position. Overlaying the local shape on top of the exterior contour (Set 3) strongly reduced the contribution of local terms, indicative of interference due to contour grouping. Local elements enclosed within and near to the exterior contour yielded local and cross-scale terms that were the strongest in magnitude, whereas local elements situated outside the exterior contour yielded weak local and cross-scale terms.

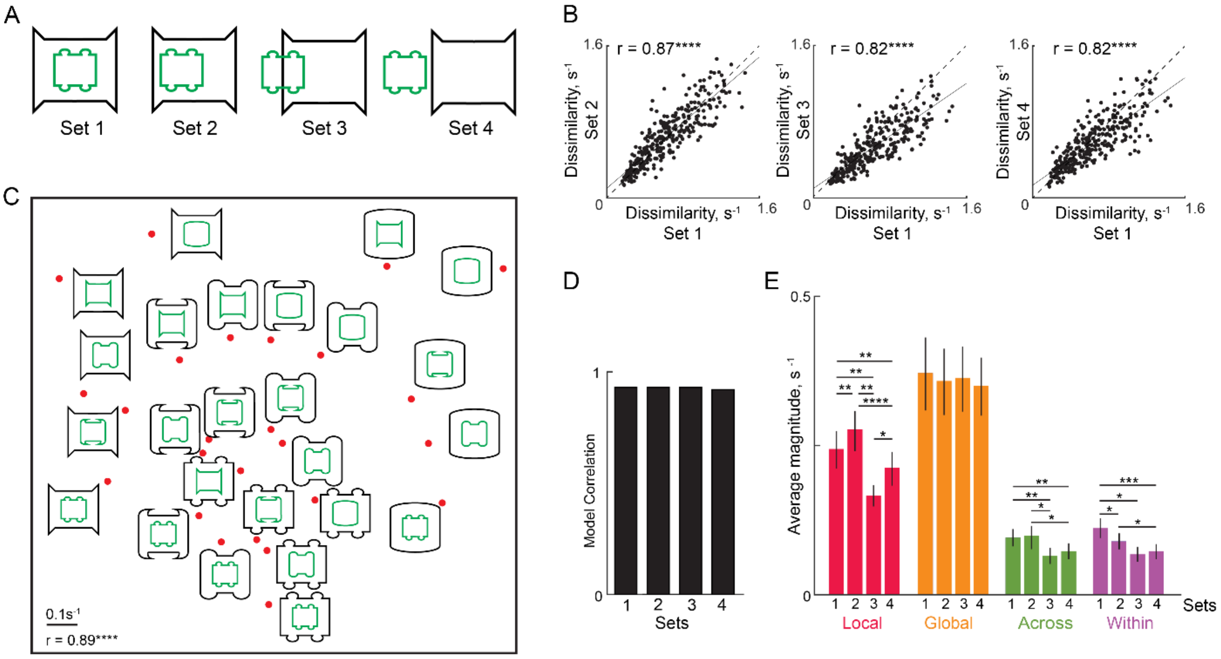


**Figure S3: Effect of local element position**

**(A)** Example stimuli from Sets 1-4, in which local elements were made larger compared to the previous experiments and were shifted along a much larger range of positions.

(B-D) Observed dissimilarity of all 300 pairs of stimuli in each set plotted against Set 1.

(E) Visualization of the underlying shape representation for the reference set (Set 1), as obtained using multidimensional scaling. All conventions are as before.

(F) Correlation between predicted and observed dissimilarities for each set.

(G) Average magnitude of model parameters for Sets 1-4. Asterisks represent statistical significance assessed using a signed-rank test: * is $p < 0.05$, ** is $p < 0.005$. All comparisons are not significant ($p > 0.05$) unless marked with an asterisk.

### SECTION S5: CHANGING ELEMENT GROUPING

Here, we examine one further influence on the shape representation, namely grouping, by creating stimuli containing identical shapes but differing in their grouping status.

#### METHODS

*Subjects.* Eight right-handed human subjects (6 male, aged 20-26 years) participated in the study. All other details are as in Experiment 1.

*Stimuli.* We created four sets of interior-exterior shapes each containing 25 stimuli. Each stimulus contained four identical interior shapes (FIgure S4A). Sets 1 & 2 consisted of stimuli in which the local elements were identical in colour (red in Set 1, green in Set 2). Sets 3 & 4 consisted of stimuli in which two local elements were green and the other two red (Set 3: green along the main diagonal, Set 4: red along the main diagonal). Thus Sets 1-2 have local elements that group by colour and shape whereas Sets 3-4 have local elements that group by shape alone. Half of the subjects performed searches involving Sets 1 & 3 and the other half performed searches involving Sets 2 & 4. In the results, we report the combined results across Sets 1 & 2 as Grouping 1 (G1) and Sets 3 & 4 as Grouping 2 (G2).

*Procedure.* Subjects performed oddball search exactly as before, with the largest item measuring 4.4°. All other details are identical to Experiment 2.

#### RESULTS

In this experiment, subjects performed oddball searches for interior-exterior stimuli that either contained local elements of identical colours (G1) or of different colours (G2). Figure S4A illustrates these two types of stimuli. Importantly because the colour and arrangement of the local elements was identical for the target and distractors, these could not serve as cues to guide visual search. Thus differences in search performance across sets can only be due to differences in grouping status.

Subjects were highly consistent in their performance across the two groups (split-half correlation between RT of odd- and even-numbered subjects: $r = 0.79$ & $0.81$ for G1 & G2 respectively, $p < 0.00005$ in all cases). Observed dissimilarity was also extremely similar across the two sets, suggesting that the underlying shape representation is qualitatively similar across grouping status (Figure S4C). To visualize the underlying shape representation, we performed a multidimensional scaling as before on the search dissimilarities of G1 pairs. The resulting plot (Figure S4B) shows that stimuli tended to group together by their exterior shape.

For both levels of grouping, the multiscale model yielded excellent fits to the data ($r = 0.93$ & $0.93$ for G1 & G2, $p < 0.00005$) indicating that systematic variations across grouping must be captured by systematic variations in model parameters. Indeed, when grouping is disrupted, global and across-object terms increased whereas local and within-object terms decreased (Figure S4D). Thus, in terms of decreasing local & within-object terms, disrupting grouping has the same effect as decreasing local element size (Figure S3H). However, disrupting grouping appears to increase across-object interactions, an effect opposite to that observed with decreased element size (Figure S3H) – this is difficult to reconcile with the other changes.

Taken together, these results show that searches for interior-exterior stimuli are explained extremely well by the multiscale model across changes in the grouping status of local elements, and that grouping tends to make local elements more salient.

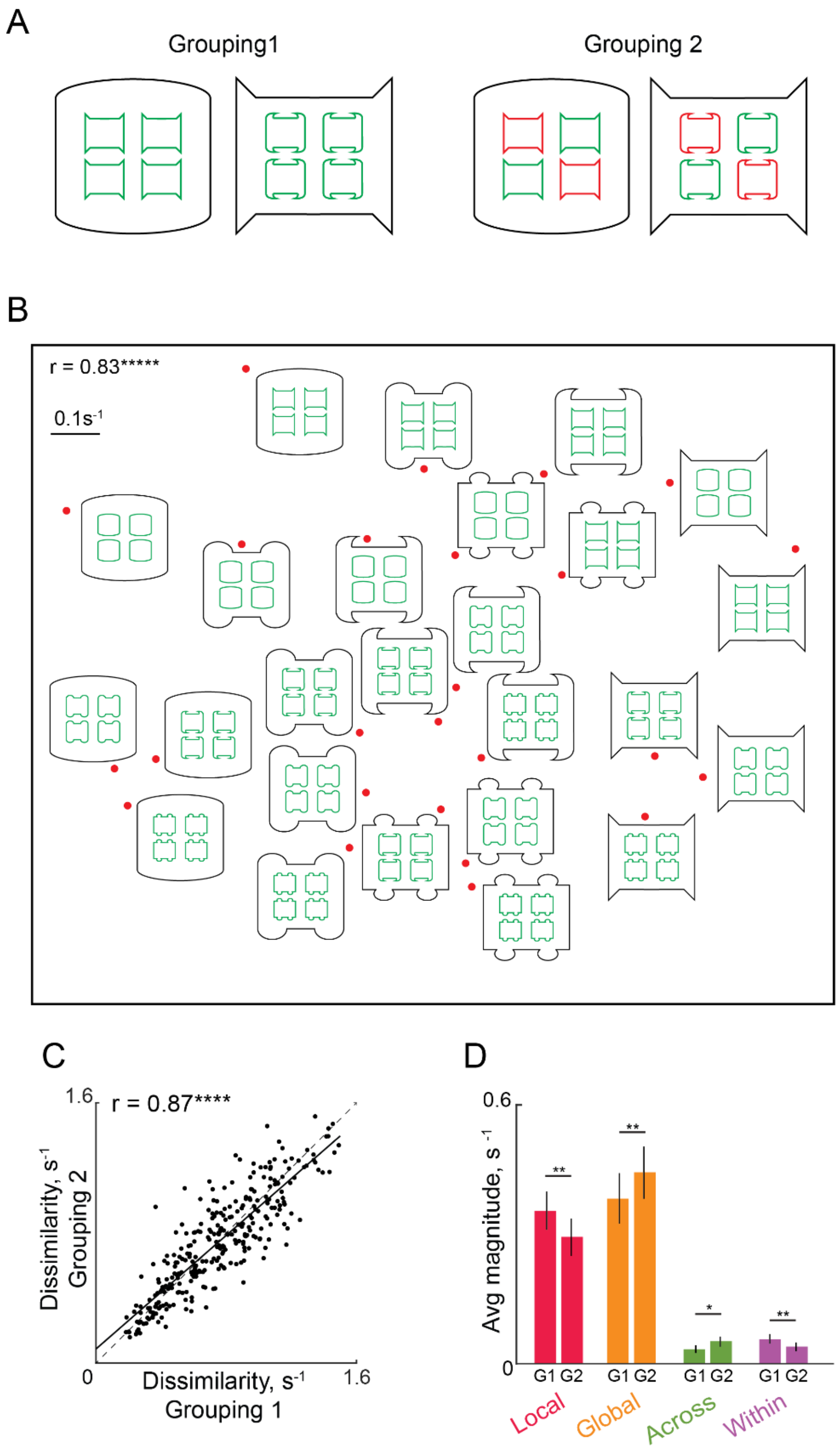


**Figure S4: Effect of element grouping on feature integration**

(A) Example stimuli from Sets 1&3 representing the two grouping levels (G1 & G2).

(B) Visualization of the underlying shape representation for the reference set (G1), as obtained using multidimensional scaling. All conventions are as before.

(C) Observed dissimilarity plotted against predicted dissimilarity for set G1.

(D) Average magnitude of model terms for sets G1 & G2. Asterisks represent statistical significance assessed using a signed-rank test: * is $p < 0.05$, ** is $p < 0.005$. All comparisons are not significant ($p > 0.05$) unless marked with an asterisk.
